# MicroRNA miR-29 controls a compensatory response to limit neuronal iron accumulation during adult life and aging

**DOI:** 10.1101/046516

**Authors:** Roberto Ripa, Luca Dolfi, Marco Terrigno, Luca Pandolfini, Valeria Arcucci, Marco Groth, Eva Tozzini Terzibasi, Mario Baumgart, Alessandro Cellerino

## Abstract

Iron is an essential metal cofactor for enzymes involved in many cellular functions such as energy generation and cell proliferation. However, excessive iron concentration leads to increased oxidative stress and toxicity. As such, iron homeostasis is strictly controlled by two RNA binding proteins known as Iron Regulatory Proteins (IRPs) that regulate at post-transcriptional level the expression of iron management genes. Despite this fine regulation, impairment of iron homeostasis occurs during aging: iron progressively accumulates in several organs and in turn, it exacerbates cellular vulnerability and tissue decay. Moreover, excessive iron accumulation within the CNS is observed in many neurodegenerative diseases. We investigated the age-dependent changes of iron homeostasis using the short lived fish Nothobranchius furzeri. Here, we show that i) both iron content and expression of microRNA family miR-29 increase during adult life and aging in the N. furzeri brain; ii) iron up-regulates miR-29 expression in fish brain and murine neurons, while in turn miR-29 targets the 3′-UTR of IREB2 mRNA, reducing iron intake; iii) Transgenic fish with knock-down of miR-29 show significant adult-onset up-regulation of IRP2 and its target TFR1 in neurons and display enhanced age-dependent accumulation of brain iron; iv) miR-29 triggers a global gene expression response that partially overlaps with that induced by aging.

Our studies indicate that miR-29 modulates intracellular iron homeostasis and is up-regulated as an adaptive response to limit excessive iron accumulation and prevent early-onset aging processes.

## Introduction

Iron is the most abundant trace element in the animal body, its abundance reflects its essential role in cell physiology. Iron is a required cofactor for carrier proteins and enzymes involved in fundamental biological process such us oxygen delivery, DNA synthesis and oxidative phosphorylation (for review see Dlouhy AC et al., 2013; Rouault TA et al., 2005). Although iron is an essential element, it is also extremely reactive and capable of producing radical oxygen species (ROS), thereby inducing oxidative damage and apoptosis (Dixon SJ et al, 2014). Intracellular iron concentration is under strict physiological control. Iron responsive elements (IRE) in mRNAs and iron responsive proteins (IRPs) constitute an iron sensing system that tightly regulates intracellular iron homeostasis coordinating iron uptake, storage, export and utilization (Zhang DL et al., 2014). IRP1 and IRP2 are RNA-binding proteins that recognize specifically IREs located in 3′- or 5′- UTRs of different protein-coding transcripts and regulate their translation (Meyron-Holtz et al., 2004; Sanchez M et al., 2011). At steady-state, IRPs positively regulate the translation of transferrin receptor (*TFR1*) and bivalent metal transporter 1 (*SLC11A2*) mRNAs (Garrick et al., 2003) and repress ferritin heavy and light-chain (*FTH, FTL*) and ferroportin (*SLC40A1*) mRNAs (Theil 1990; Abboud and Haile, 2000; McKie et al., 2000) thereby promoting iron uptake and availability. However, in iron-repleted conditions, F-Box And Leucine-Rich Repeat Protein 5 (FBLX5), stabilized by iron, starts to accumulate and interacts with IRP2 mediating its ubiquitination and degradation (Salahudeen A et al., 2009; Vashisht A et al., 2009) instead IRP1, via conformational change, loses the RNA binding ability and acquires cytosolic aconitase activity (Haile DJ et al., 1992). Thus, IRPs inactivation positively regulates *FTH, FTL* and *SLC40A1* and represses *TFR1* and *SLC11A2* mRNA translation reducing iron uptake and promoting iron storage and export. While IPR1 is mostly expressed in liver, spleen, blood and kidney, IRP2 is strongly expressed in the central nervous system (CNS). IRP2-deficienct mice display severe iron metabolism misregulation in the CNS and develop movement disorders characterized by ataxia, bradykinesia and tremor (LaVaute T et al., 2001). Instead, transgenic overexpression of IRP2 in mice increases neuronal iron up-take and leads to mitochondrial oxidative insults to the CNS and a subsequent degeneration of dopaminergic neurons (Asano T et al., 2015).

Despite the presence of a sophisticated homeostatic regulation, during aging iron accumulates in multiple tissues and this likely contributes to physiological decline (Bartzokis G et al., 1997; Zecca L et al., 2004). Dietary iron supplementation was shown to induce protein insolubility and significantly reduce *C. elegans* lifespan (Klang IM et al., 2014). In particular, the age-dependent iron accumulation in the central nervous system (CNS) was linked to cognitive dysfunctions and memory deficit in humans (Penke L et al., 2012). Furthermore, marked iron accumulation is a hallmark of neurodegenerative diseases like Alzheimer Parkinson and multiple sclerosis, increasing ROS production thereby exacerbating oxidative damage and cell death (Smith et al., 1997; for review see: Crespo et al., 2014; Wong et al 2014; Oshiro et al., 2011). In the context of Alzheimer’s disease, iron accumulation can promote the formation of both Aβ plaques and tau-tangles (Mantyh et al., 1993; Yamamoto et al., 2002). Furthermore, Amyloid precursor protein (APP) and α-Synuclein expression are modulated at post-transcriptional level by IRE-IRPs system and recently APP was shown to be required for iron export (Duce et al., 2010; Febbraro et al., 2012; Friedlich et al., 2012). Therefore, dysregulation of iron can be considered as one of the key pathophysiological mechanisms of aging and age-related neurodegenerative diseases.

MicroRNAs are small non-coding RNAs that form a ribonucleic complex with argonaute proteins. This complex negatively regulates expression of protein-coding genes at a post-transcriptional level by binding mRNAs due to sequence complementarity and inducing mRNA degradation and translational inhibition. MiR-29 family members (miR-29a, miR-29b and miR-29c) are produced in mammals from two intergenic *loci* and are one among the most expressed microRNA families (Landgraf et al. 2007). They are enriched in the CNS, are expressed both in neuronal and glial cells and their binding sites are highly overrepresented in brain mRNAs co-immunprecipitated with argonaute complex (Boudreau et al., 2014). During brain development of rodents and primates, miR-29 is expressed at low levels and its expression dramatically increases during late postnatal development and continues to increase during entire adult life and aging in humans and non-human primates (Somel et al., 2010; Podolska et al., 2011; Fenn et al., 2013). In addition, age-dependent up-regulation of miR-29 was observed also in other organs such us heart, lung, liver, kidney of mice, and aorta of both mice and humans (Boon et al., 2011; Takahashi et al., 2012, Ugalde et al., 2011). Therefore, up-regulation of miR-29 seems to be a conserved hallmark of aging.

Mice lacking only the miR-29b/a-1 *locus* (therefore with residual miR-29 activity) reach adulthood, display ataxic phenotype and a mild loss of Purkinje cells in the cerebellum and die around 9 months of age (Papadopoulou et al., 2015). A similar cerebellar phenotype is induced acutely by miR-29 antagomiR (Roshan et al., 2014). On the contrary, mice with a targeted deletion of all MiR-29 *loci* are born without evident abnormalities, but rapidly accumulate defects during post-natal development and die within 6 weeks (Cushing et al., 2015). Interestingly, this microRNA family was found to target Beta-Site APP-Cleaving Enzyme (*BACE1*) mRNA and to be downregulated in sporadic Alzheimer’s disease (Hébert et al., 2008). Moreover, up-regulation of miR-29 protects neurons against apoptosis during neuronal maturation, forebrain cerebral ischemia and stroke by targeting pro-apoptotic members of the BCL-2 family (Kole et al., 2011; Ouyang et al., 2013; Khanna et al., 2013). Overall, these data strongly suggest a conserved key role for miR-29 family in protecting neurons from aging-induced damages.

Here, we investigated the role of miR-29 during adult life in the short-lived killifish *Nothobranchius furzeri*, an emerging vertebrate model species for aging research, characterized by an exceptionally short lifespan of 4-12 months, age-dependent cognitive decline and expression of age-related phenotypes at the molecular, cellular and integrated level (Valenzano et al., 2006a; Valenzano et al., 2006b; Di Cicco et al., 2011; Terzibasi et al., 2012; Wendler et al., 2015; for a review see Cellerino et al., 2015). Moreover, transgenesis was established, a reference genome sequence is available and extensive data on age-dependent gene regulation are available (Valenzano et al., 2011; Baumgart et al., 2014; Valenzano et al., 2015; Reichwald et al., 2015: Baumgart et al., 2016). We previously demonstrated that expression of miR-29 increases during aging in *N. furzeri* (Baumgart et al., 2012). Here, we investigate the physiological relevance of miR-29 *in vivo* experimentally by generating genetically-modified fish lines expressing a competitive inhibitor for miR-29 (sponge) in the neurons. Further, we show that miR-29 targets IRP2 and test the hypothesis that miR-29 directly regulates brain iron homeostasis and opposes age-dependent iron accumulation and the resulting damages.

## Results

### Iron accumulates in *Nothobranchius furzeri* brain

We investigated age-dependent brain iron accumulation in *N. furzeri* MZM 04/10pl, a strain with a median lifespan of ~ 30 weeks (Terzibasi *et al*., 2008, 2009; Baumgart *et al*., 2014). Brain non-heme iron concentration was quantified using a colorimetric assay (Rebouche *et al*., 2003) in fish of five age groups: 5, 12, 20, 28 and 39 weeks. These age steps correspond to sexual maturity, young adult, mature, median lifespan and very old stages. We found that, like in mammals (Hallgren *et al*., 1958; Bartzokis *et al*., 1997), iron content increases over time. From 5 to 39 weeks of age, iron amount increases almost 10-fold in animals of both sexes (two-way ANOVA age: P<0,0001, gender: P=0,75, fig. 1A). This datum was corroborated by histochemical Pearl’s staining with DAB intensification on brain sections. Increased labelling was apparent in the brains of older fish, particularly in Purkinje cell layer of the cerebellum (Fig. 1B). We also investigated the expression of genes coding for key proteins of iron metabolism by interrogating a public dataset of genome-wide age-dependent transcript regulation in the *N. furzeri* brain at the same five age steps analyzed here (Baumgart et al., 2014). We observed that expression of the transferrin receptor (*TFR1A*) gene and the solute carrier transporter 11a2 (*SLC11A2*) gene, directly involved in intracellular iron delivery, significantly declines as a function of age (ANOVA for linear trend R=0,5009, P<0,0001; R=0,3892, P<0,01, respectively Fig. 1C), with the largest difference observed between 5 weeks and 12 weeks of age. On the other hand, expression of the genes coding for ferritin heavy chain (*FTH1A*) and Ferroportin (*SLC40A1*), required for iron storage and discharge, are stable (ANOVA for trend R=0,067, P=0.11; R=0.078; P=0,11, respectively Fig. 1C). Since all of these genes are controlled by IRE-IRPS system, we investigated whether expression of *IREB2* and *ACO1* mRNAs is age-dependent and did not detect significant regulation during aging (ANOVA for trend R=0,141, P=0,069; R=0.045, P=0,33 Fig. 1D). *IREB2* mRNA is higher-expressed in brain as compared to *ACO1* mRNA, while the opposite is observed in the liver (Fig. S1A). These same differences in the relative expression levels of *IREB2* and *ACO1* between brain and liver were previously observed in the mouse (Meyron-Holtz et al., 2004) and suggest that *IREB2* is the principal regulator of iron homeostasis in the brain. In mammals, iron regulates *IREB2* expression mainly at the post-translational level by iron-dependent proteasomal degradation of IRP2. As expected, Western blot analysis revealed that IRP2 decreases with age (ANOVA for linear trend R=0.7199, P=0.0002, fig. 1E). Since IRP2 is negatively regulated by FBXL5 we evaluated also its expression at mRNA and protein level but curiously in both cases we didn’t found any significant variation over time (Fig. S1B-C). These data suggest that iron progressively accumulates over time and IRP2 undergoes toward an age-dependent downregulation not FBXL5 mediated.

**Figure 1.**
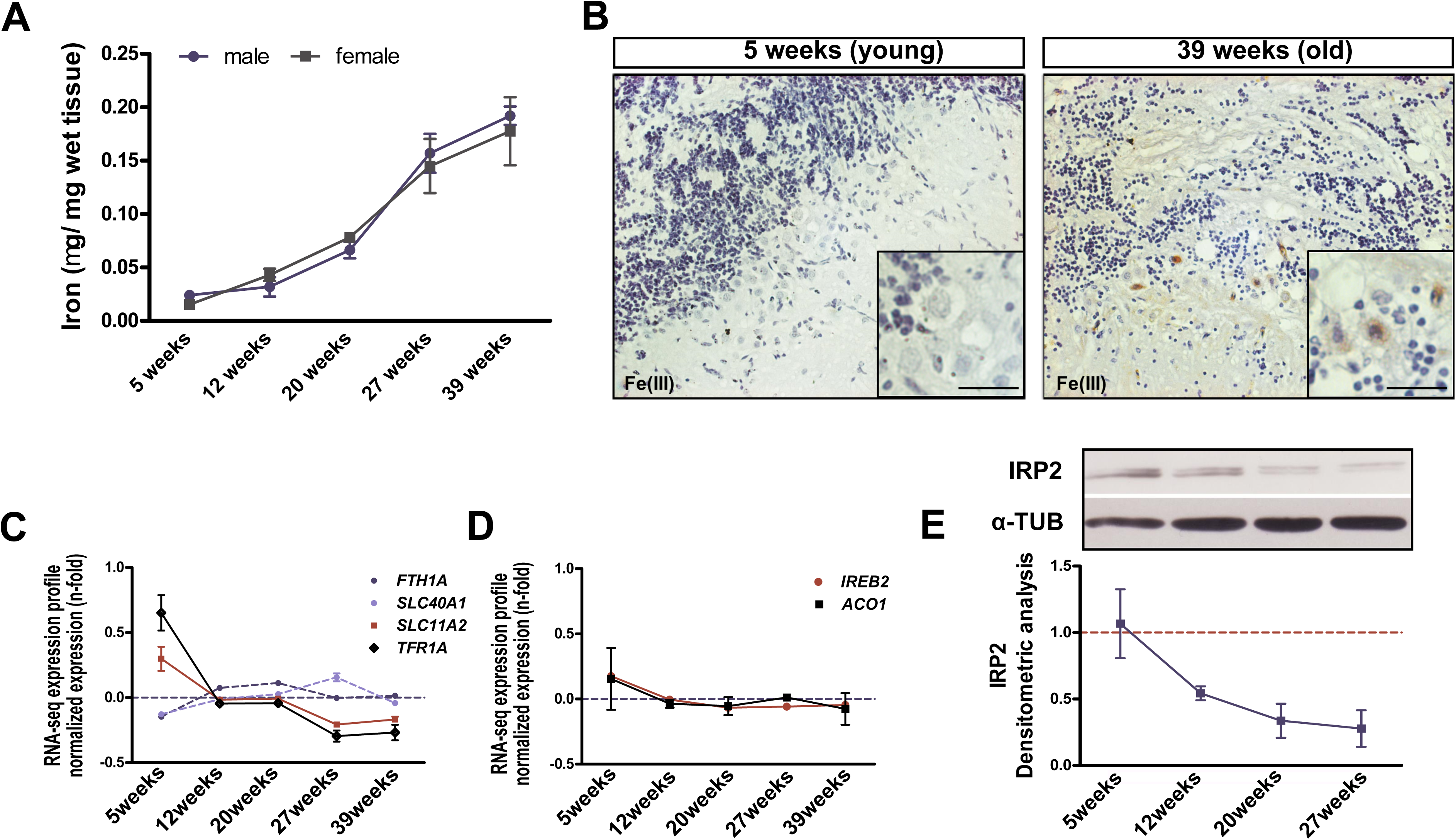
<u>Age dependent brain iron accumulation in *Nothobranchius furzeri*</u> **A)** Brain non-heme iron content (μg/g wet tissue) in fish of different ages. Blue and black lines represent males and females, respectively. Iron increases with age, (P<0.0001, two-way ANOVA), and no differences were observed between the sexes (P=0.75). **B)** DAB-enhanced Perl’s staining of the cerebellum of young and old fish. The black arrows point to labelled Purkinje cells. The inset in the bottom right corner of each picture shows a magnification of the Purkinje cells bodies. **C-D)** Age-dependent regulation of the transcripts coding for key genes of iron metabolism, from Baumgart et al., (2014). Expression values (in RPMKs) for each age were centered and scaled to the mean. **E)** Representative Western blot of IRP2 in brain extracts and densitometric analysis. α-TUBULIN was used as loading control. For C, D mean ± starndard errors of means is reported and the significance of age-dependent modulation was assessed using 1 way ANOVA with post-test for trend.

### MiR-29 family targets *IREB2* mRNA

MicroRNA-29 is up-regulated with age in multiple tissues of *N. furzeri* including brain (Baumgart et al., 2012). We noticed the presence of an 8 nt seed sequence in the 3′UTR of *IREB2* mRNA that is evolutionary conserved in vertebrates (Fig. 2A) and both miRanda (Betel D et al., 2010) and TargetScan (Agarwal et al., 2015) retrieved *IREB2* mRNA as a target for miR-29 both in fishes and mammals. In addition, StarBase provides support for a physical interaction of miR-29 and *IREB2* 3′-UTR in human and mouse (Li J *et al*., 2014). We then investigated the transcriptional control of miR-29. The *N. furzeri* genome contains three loci for miR-29 family members but one is expressed at much higher levels, (a detailed analysis of *N. furzeri* miRNA repertoire is in preparation and will be published elsewhere). While IRP2 is down-regulated with age, transcription of the miR-29 *locus* is up-regulated more than 15 times with age (Fig. 2B). In addition, we investigated transcriptional control of miR-29 in the mouse brain since it was already shown that mature miR-29 dramatically increases during the first two months of postnatal life (Hebert S et al., 2008; Li H et al., 2013). Using RT-qPCR, we quantified the expression of both miR-29 primary transcripts in postnatal day 0 (P0) and P60 mouse cerebral cortex. As expected, both increased more than 30 times (miR-29b/a P<0,001, miR-29b/c P<0,001, fig. 2C). Then, we assessed the expression of iron management genes such as *IREB2*, *TFRC*, and *DMT1*: all were found to be significantly downregulated (P<0,01, P<0,05, P<0,05, respectively Fig. 2D). It also relevant to note that mice lacking one locus for miR-29 show up-regulation of *Ireb2* (Papadopulous et al., 2015). These data suggest that both in fish and mouse miR-29 is involved in the age-dependent down-regulation of IRP2. We further used *in situ* hybridization to define miR-29 expression domains in the adult *N. furzeri* brain. We found this primary transcript intensely expressed in the periventricular gray zone (PGZ) of the optic tectum (fig. 2E), and in the granular cell layer (GCL) of cerebellum (fig. 2F), where it was expressed by neurons, as showed by its co-localization with the neuronal marker HuC/D. To demonstrate a direct interaction between miR-29 family members and *IREB2* mRNA in fish, we isolated the 3′UTRs of *IREB2* from zebrafish and *N. furzeri* cDNA and fused them with the eGFP coding sequence to generate a reporter system. 100μg of each reporter mRNA were co-injected in fertilized zebrafish embryos with the same amount of control mRNA encoding for the red fluorescence protein (RFP) and 100μg of dre-miR-29a or dre-miR-29b mimic. Control embryos were injected with the same mixture with omission of the miRNA mimics. At 24hpf, injected embryos were trypsinized and analyzed by cytofluorimetry to assess the eGFP/RFP ratio (fig. 2G). For both 3′UTRs, the normalized eGFP fluorescence was significantly reduced in presence of miR-29 mimics (35%, P<0,05 T-test and 40% P<0,01, T-test for *N. furzeri* and zebrafish 3′UTR respectively, fig. 2H-I). For both constructs, a mutation of 4 nucleotides in the putative target sequence of the 3′-UTRs abolished repression (fig. 2H-I). In addition, we performed luciferase assay in HEK293T cells to demonstrate direct targeting of *IREB2* 3′UTR by miR-29 in mammals. MiR-29 decreased luciferase activity of a reporter containing mouse *IREB2* 3′UTR (P<0,01 T-test;) and repression was abrogated by mutation of the putative binding site (fig. 2J).

**Figure 2.**
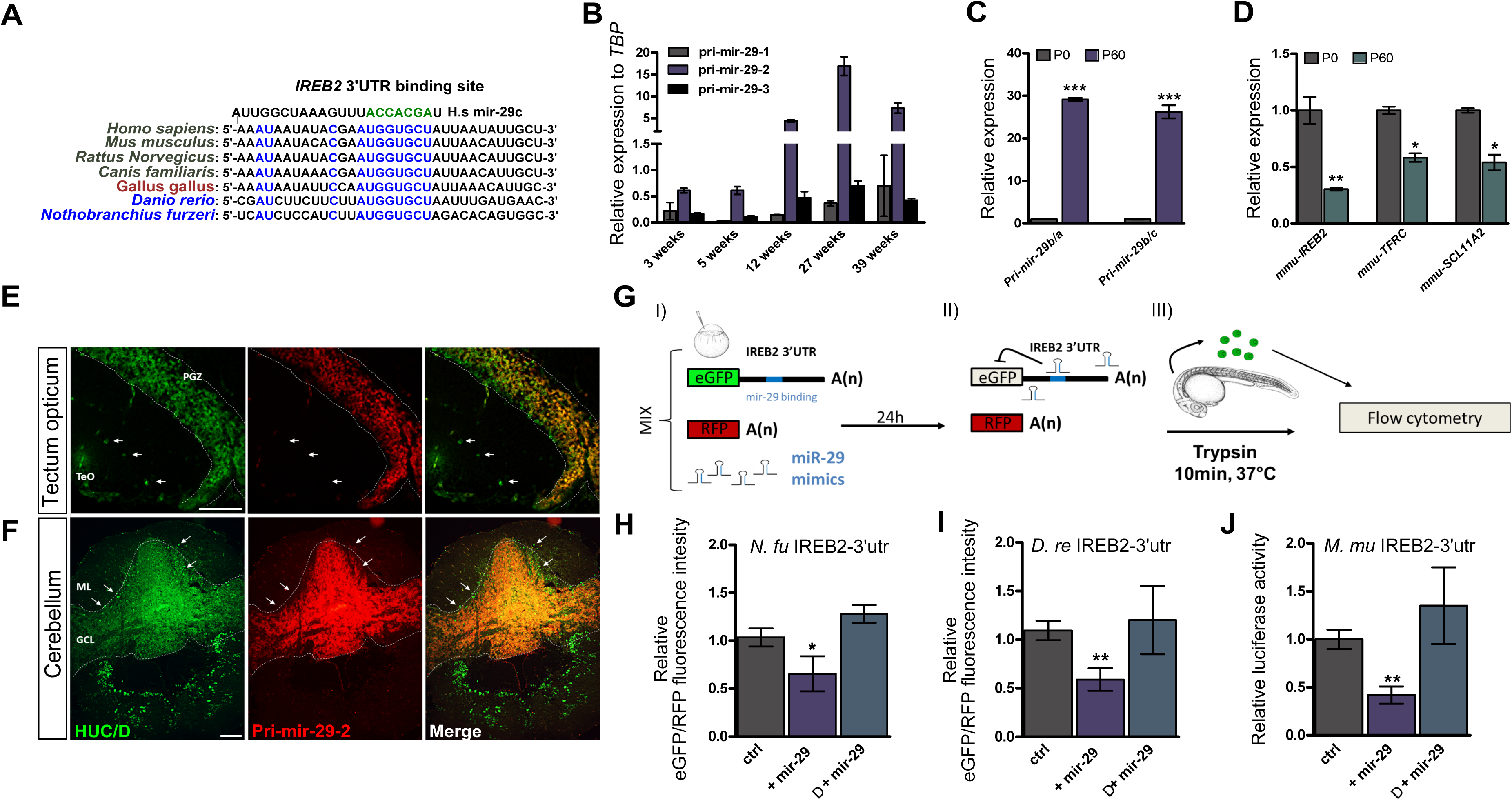
<u>MiR-29 family targets *IREB2* mRNA</u> **A)** Presence of a putative binding site for miR-29 family in the *IREB2* mRNA 3′-UTR sequence in several vertebrate species (*H. sapiens*: ENSG00000136381; *M. musculus*: ENSMUSG00000032293; *R. norvegicus*: ENSRNOG00000013271; *C. familiaris*: ENSCAFG00000001766; *G. gallus*: ENSGALG00000003171; *D. rerio*: ENSDARG00000021466; *N. furzeri*: Nofu_GRZ_cDNA_3_0193494), mammalian sequences are in black, birds in purple and teleost fish in blue. Perfect match to the position 2-8 of the miR-29 seed sequence is highlighted in green and is present in all vertebrate sequences shown. **B)** Age-dependent expression of miR-29 primary transcripts (Pri-miR-29-1, 2, 3) in the brain of *N. furzeri*. The relative expression was evaluated by RT-qPCR, data were normalized on TATA binding protein (TBP), pri-miR-29-2 results much more expressed than the other clusters and shows a clear age-dependent up-regulation (1 way ANOVA with post-test for trend: R=0,5285 P<0,0001, n=4 biological replicates for age group). **C)** Quantification of pri-miR-29 cluster (pri-miR-29b-a; pri-miR-29b-c) in post-natal day 0 and 60 mouse brain. Both primary transcript dramatically increase during post-natal development (P<0,001 and P<0,001 respectively). **D)** Relative expression of iron management genes *IREB2* (P<0,01) *TFRC* (P<0,05), *SLC11A2* (P<0,05) in post-natal day 0 and 60 mouse brain. Both for C and D statistical significance was calculated by Mann-Whitney’s U-test, n=6 (pO brain) and n=7 (p60 brain) biological replicates. **E-F)** Pri-miR-29-2 expression pattern in *N. furzeri* brain. Optic tectum magnification (E) shows pri-miR-29-2 *in situ* hybridization signal (red) and HuC/D expression (green). Pri-miR-29-2 shows a nuclear staining and a co-localization with neuronal marker HuC/D in all periventricular gray zone (PGZ), white arrows show neurons in the optic tectum (TeO) negative for pri-miR-29-2. Scale bar 50μm. Cerebellum overview picture **(F)** shows a clear and strong expression of pri-miR-29-2 just in the granular cell layer (GCL), it is instead is absent in the Purkinje cell (white arrow) and molecular layer (ML). Scale bar 200μm. **G)** Scheme of the reporter constructs and assay for miR-29 activity. Green fluorescent protein (GFP) mRNA fused with the 30 UTR of *IREB2* is injected in one-cell stage zebrafish embryos with or without the miRNA of interest. Red fluorescent protein (RFP) mRNA is injected as a loading control (i). Binding of miR-29 to the reporter mRNA causes repression of GFP signal (ii). Embryos were trypsinized and cells relative fluorescence was read by flow cytofluorimeter (iii). H-l) Expression of GFP assessed by cytofluorometric analysis. Fusion with *IREB2* 3′UTR of *N. furzeri* **(H)** and D. rerio **(I)**. The white bars indicate the baseline fluorescence intensity of the construct shown in (G) in the absence of miR-29 mimic. The middle black bars indicate the fluorescence in the presence of miR-29 mimic and the right black bars the fluorescence of a construct (A) where the putative binding site for miR-29 in the *IREB2* 3′-UTR was mutated to destroy complementarity. Statistical significance of fluorescence difference between baseline and co-injection with miR-29 mimics was evaluated by Student’s t-test (*, P<0,05; **, P<0,01). **J)** Dual luciferase assays of 293T cells co-transfected with firefly luciferase construct containing the wild-type or the mutant target sites of mmu-*IREB2* 30-UTR along with the miRNA expression plasmid (pcs2+: CMV:RFP-miR-29b/c precursor-polyA-tail) or the empty vector (pcs2+: CMV:RFP-polyA-tail). Histograms show normalized mean values of the relative luciferase activity of miRNA expression plasmid-transfected cells with respect to empty vector transfected cells, from 4 tests of two independent transfections. Values are normalized to the mean of control plasmid transfected cells. Bars represent meant standard deviation (**, P<0,01; T-test)

### Iron overload induces miR-29 in neurons

Given the regulation of *IREB2* by miR-29, a logical question to ask was the regulation of miR-29 by iron. First, we studied stem-cells derived murine telencephalic neurons *in vitro* (Bertacchi M *et al*., 2014). Incubation of murine neurons with either Fe(II) or Fe(III)-dextran induced a dose-dependent up-regulation of miR-29 (Fig 3A-B). We then induced acute iron overload in adult fish by parental injection of 350μg per gram of weight of Fe-dextran and monitored brain iron concentration and miR-29 expression up to three days post-injection (p.i.). Iron significantly accumulated in the brain 4 hours p.i. (the first time point investigated) reaching a maximum at 8-12 hours p.i. to slowly decrease (fig. 3C). We observed a delayed and significant increase of both miR-29 primary transcript 2 and miR-29a mature form (fig. 3D), following the injection, starting at 24 h p.i. and reaching its maximum at 48h p.i. (pri-miR-29 48h: P<0,01; 72h: P<0,05; miR-29a 48h P<0,05; 72h P<0,05, one way ANOVA; fig. 3C). In addition, iron overload caused up-regulation of miR-29 also in liver and muscle (fig. S3A), suggesting that this regulation is not brain specific, but it takes place at systemic level. We then asked whether physiological steady-state levels of iron could influence miR-29 expression. We injected i.p 30μg/g of the iron chelant deferoxamine (DFO). At 24 hours post-injection brain iron amount was reduced by 25% (P<0,05, fig. S3A), we did not aim for stronger reduction of iron since this can have serious negative consequences on physiology. We monitored pri-miR-29-2 expression at 24h, 48h and 72h after injection and did not detect any differences (fig. 3E), suggesting that physiological levels of iron do not regulate miR-29. Induction of miR-29 might be a direct response to iron accumulation or a secondary response to iron-induced oxidative stress. To distinguish between these two possibilities, we replicated the iron-overload experiment and, after 4 hours, we administered either DFO, and we assessed miR-29 expression at 48h after iron injection. DFO treatment significantly reduced pri-miR-29 up-regulation. (P<0,05, fig. 3F). Our results suggest that intracellular iron increase directly up-regulates miR-29.

**Figure 3.**
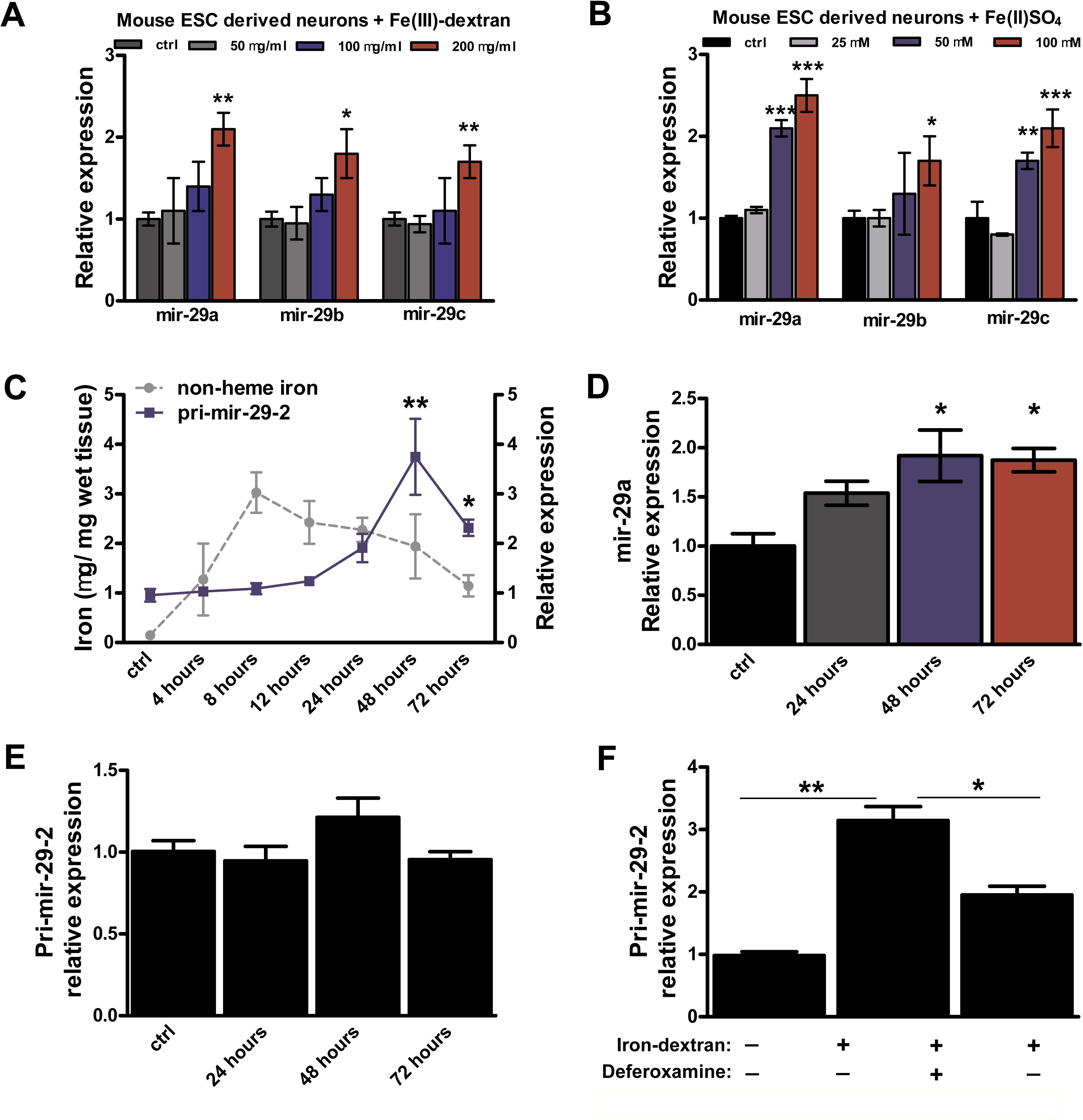
<u>Iron overload induces miR-29 up-regulation in neurons and brain</u> **A-B)** Modulation of miR-29 family members in murine neurons derived from mESCs incubated respectively with: **A)** 0, 50μg/ml, 100μg/ml, 200μg/ml of Fe(III)-dextran or with **B)** 0, 25μM, 50μM, 100μM of Fe(II)SO4 for 72 hours. Statistical significance between control and iron-treated cells was evaluated by one-way ANOVA with post-hoc Tukey’s test (* = P<0,05; ** = P<0,01; 046516 = P<0,001), n=3 independent replicates. **C)** Time course of non-heme iron amount (quantified by colorimetric analysis) and pri-miR-29-2 expression level in fish brain (quantified by RT-qPCR) following intraperitoneal (i.p.) injection of 350 μg/g of iron dextran, control animals were injected with saline solution. Grey line represents iron amount, blue line represents pri-miR-29-2 relative expression level (setting the baseline to 1). Statistical significance for pri-microRNA expression was calculated by one-way ANOVA with post-hoc Tukey’s test (*, P<0,05; **, P<0,01), n=4 biological replicates for each point. **D)** Expression of mature A/./u-miR-29a at 48 hours after iron injection quantified by RT-qPCR, U6 was used as normalization control. **E)** Time course of pri-miR-29-2 expression level (quantified by RT-qPCR) following i.p. injection of 30 μg/g of deferoxamine (DFO), n=4 biological replicates for each time point. **F)** Modulation of pri-miR-29-2 expression in fish brain (quantified by RT-qPCR) after iron overload and in combination with administration of 30 μg/g of DFO, control animals were injected with saline solution, Mann-Whitney’s U-test (*, P<0,05); n=5 for each experimental point. For all the graphs, mean ± standard errors of means are reported.

### Genetic repression of miR-29 function

To investigate the physiological relevance of miR-29 in the regulation of iron homeostasis during adult life, we generated transgenic zebrafish and *N. furzeri* expressing a competitive inhibitor of miR-29 designed by fusing to the eGFP a 3′-UTR with seven repetitions of a miR-29 binding site (miR-29-sponge) (fig. 4A, on the top). As a first step, we tested whether miR-29-sponge transcript was directly targeted by miR-29 family members by co-injecting them with the control RFP mRNA (as described above for miR-29 mimics) measuring the eGFP/RFP ratio by fluorocytometry. A strong repression of eGFP signal was induced by miR-29 mimics (fig 4B). Then, we generated a transgenic zebrafish line by placing miR-29-sponge under the control of 5,3 kb of *D. rerio* actin beta 1 promoter (tg:actb2-eGFP-sponge-29) (fig. 4A). We then used the F1 generation to assess the functionality of the construct *in vivo*. For this purpose, we took advantage of the extremely low expression of miR-29 during the first two days of embryonic development (Chen PY *et al*., 2005) and injected increasing doses of miR-29a or miR-29b mimics, (~100μg, ~150μg and ~200μg) both in wild-type and in tg:actb2-eGFP-sponge-29. We found that both mimics induced similar effects and 200 μg of mimic induced morphological defects like brachyury, microcephaly and microphthalmia. The percentage of defective embryos was much lower in tg:actb2-eGFP-sponge-29 (fig. 4C-D). Recent work showed that at steady-state microRNAs regulate their targets more through mRNA destabilization than by translation repression (Eichhorn SW *et al*., 2014), so we used injected embryos with lower mimic dosages (100μg and 150μg by which did not display overt morphological aberration) in order to evaluate the expression of well-known miR-29 targets. We chose as target *ELNA1* (elastin) and *COL11A1A* (collagen type XI, alpha 1a), because these are genes highly expressed in embryos bearing in their 3′-UTR 4 and 2 predicted target binding sites, respectively (Fig. S4A) and quantified their expression through RT-qPCR. We observed a dose-dependent down-regulation of these transcripts in wild-type, but not in tg:actb2-eGFP-sponge-29 injected embryos. All these data supported the evidence that genetically modified fish with stable sponge overexpression could counteract miR-29 effects. Finally, we measured *IREB2* mRNA but we didn’t detect down-regulation after miR-29 mimic injection (fig. 4G), however we observed downregulation of *TFR1A* mRNA in wild type but not tg:actb2-eGFP-sponge-29 injected embryos (fig. 4H). These data suggest that, in fish, miR-29 influences IRP2 translation but not *IREB2* mRNA stability.

**Figure 4.**
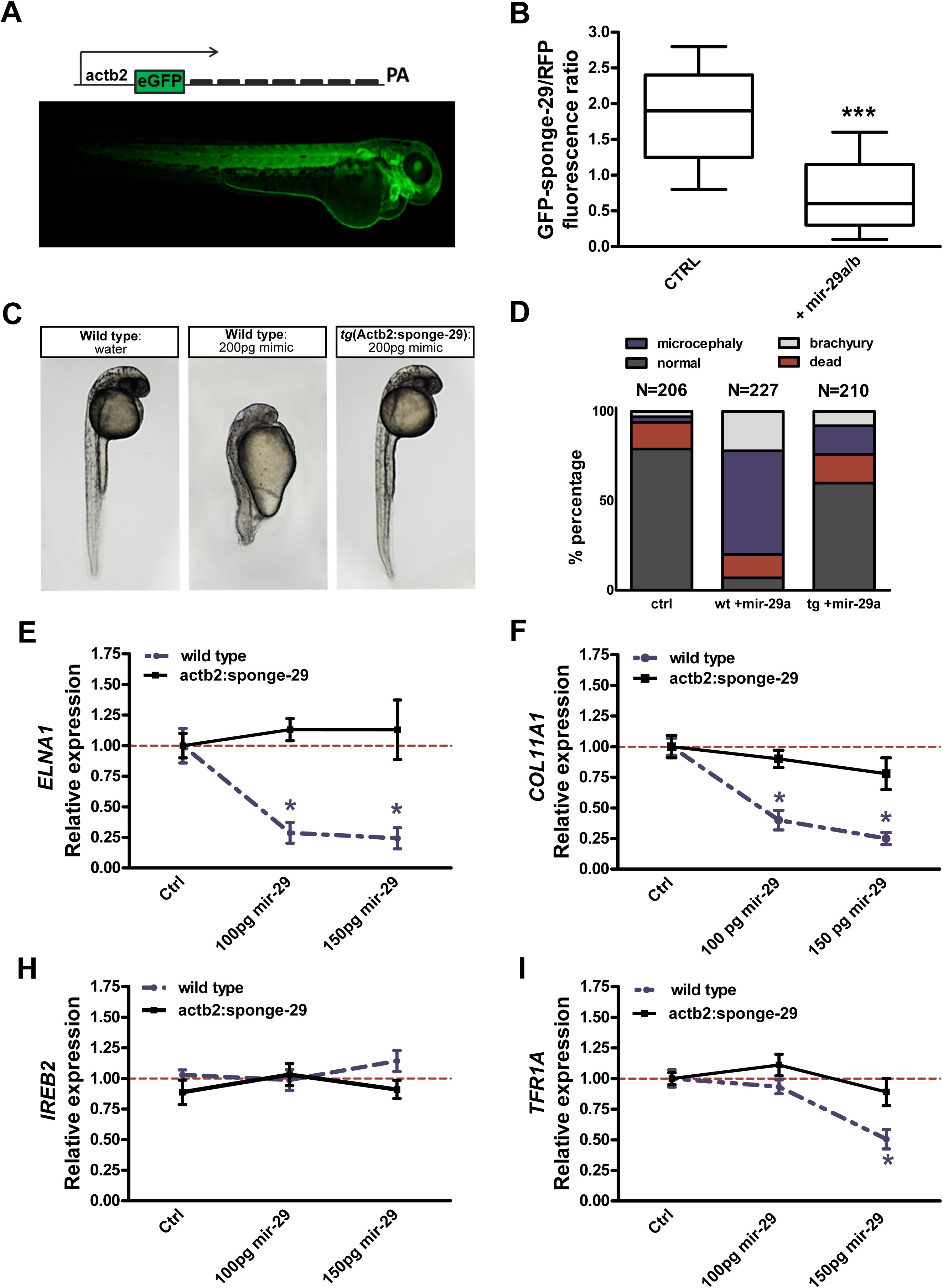
<u>Genetic repression of miR-29 function by sponge technology</u> **A)** Transgenic sponge-29 zebrafish are generated by Tol2-mediated transgenesis, expression cassette consist of 5,3kb of zebrafish actb2 promoter, eGFP fused to a synthetic 3′UTR containing 7 repetition of miR-29 binding site and a SV40 late poly-A tail. The image shows a F1 embryo (72 hours post fertilization) and the ubiquitous expression of eGFP-sponge throughout the body. **B)** Expression of eGFP measured by cytofluorimetry analysis. eGFP mRNA fused with sponge-3′UTR was injected in zebrafish embryos along with or without dre-miR-29 mimics. RFP mRNA was injected as loading control. MiR-29 mimics strongly reduced eGFP-sponge signal (046516, P<0,001, T-test). Whisker plots indicate the 10%, 25%, median, 75% and 90% ranges. **C-D)** Wild-type and F1 transgenic zebrafish were injected with 200μg of miR-29a or miR-29b mimics at the one cell stage, control embryos were injected just with water and red-phenol. Picture (C) shows representative control embryos, wild-type embryos +200μg mimics, transgenic embryos + 200μg mimics at 24hpf. Stacker bar chart (D) represents the percentages of embryos with different phenotypes (normal, death, microcephaly and brachyury) in the three conditions. **E-H)** Expression level at 24 hours post-fertilization of COL11A1, ELNA, *IREB2* and *TFR1A* upon miR-29 mimics injections. One-stage wild-type and transgenic embryos were injected with different doses (100 and 150 μg) of microRNA mimics and the expression level determined by RT-qPCR. Statistical significance was assessed by one-way ANOVA * = P<0,05. Error bars indicate standard errors of means.

### MiR-29 deficiency induces iron imbalance during adult life

Once established that miR-29 sponge counteracts the endogenous miR-29 activity *in vivo* and could perturb regulation of iron-management genes, we investigated whether and how miR-29 disruption affects iron homeostasis in neuronal cells during adult life. To this end, we isolated from zebrafish genomic DNA a 3.1 Kb neuronal specific promoter Dre-kif5a (kinesin 5a), in order to limit miR-29 loss of function to mature neurons. Kif5a promoter fragment was sufficient to maintain the expression of eGFP-sponge-29 throughout life both in zebrafish and *N. furzeri*. Since zebrafish life expectancy is of about five years (Gerhard GS *et al*., 2002) we decided to generate a stable line in *N. furzeri* in order to investigate more rapidly the consequences of miR-29 depletion during adult life. *N. furzeri* F1 fish exhibited a stable expression over time (fig. 5A-B) and a double labeling with eGFP and HuC/D proved the neuronal specificity of the expression pattern (fig. S5A). We did not observe any macroscopic defects, but a reduced fertility, and a reduced post hatch survival that limited the number of animals that could be analyzed. It should be noted that a full knock-out of miR-29 is postnatally lethal in the mouse, so reduced viability is expected and the fact that some transgenic fish reached adulthood could be due to the incomplete antagonism of miR-29 by the sponge construct. Following the initial hypothesis that neuronal miR-29 deficiency could lead to a dysregulation of iron homeostasis, we first measured by Western blot the expression of IRP2 and *TFR1A* in young adult (12 weeks old) wild type and transgenic fish. As expected, both proteins were found up-regulated as compared to control animals (fig. 5C). In addition, we assessed IRP2 level also in 5 months old zebrafish kif5a:sponge-29 line brain confirming its up-regulation (fig. S5B). Confirmed that mir-29 deficiency increased the expression or IRP2, we measured IRP2 levels at 5, 12, 20, 27 weeks of age both in transgenic animals and wild type. We found a comparable expression in young age (5 weeks), but when fish grow old the relative protein level remain largely deregulated in genetically modified fish, indicating that miR-29 significantly influences IRP2 expression during adult life (ANOVA for linear trend, wild-type: R=0,7199, P=0,0002, Sponge-29: R=0,2871, P=0,0691, fig. 5D-E). Therefore we measured both iron-management genes expression and iron content in young adult 12 weeks old wild type and kif5a:sponge-29 fish. We quantified by RTq-PCR the expression of *IREB2, TFR1A, FTH1A, SLC401A, SLC11A2*, but we found a significant difference only for *TFR1A*, up-regulated in the kif5a:sponge-29 animals (p<0,05; fig. 5F). This is consistent with the notion that iron homeostasis in neurons is regulated mainly by the ortholog of *TFR1A*, *TFRC* since they intake iron in the form of transferrin-bound iron released by the astrocytes (Rouault, 2013) and variations of TRF1A expression would have the largest consequences on neuronal iron homeostasis. Finally, we quantified non-heme iron content and this was significantly increased as compared to wild type (P<0,05, fig. 5G), suggesting that miR-29 deficiency accelerates iron in-take over time.

**Figure 5.**
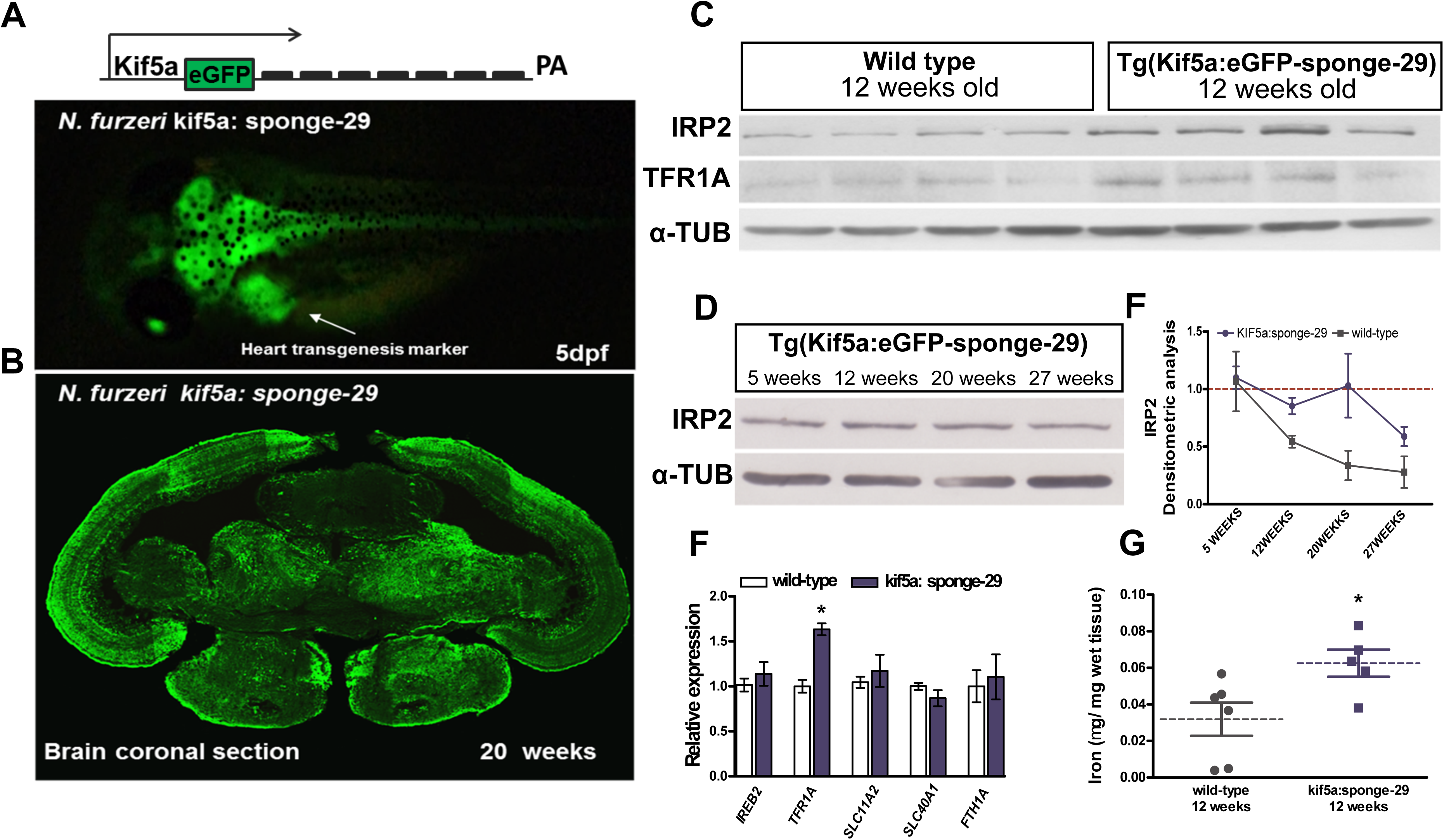
<u>Genetic repression of miR-29 chronically affects iron homeostasis during adult life</u> **A)** Schematic representation of the kif5a:eGFP-sponge-29 expression cassette and F1 *N. furzeri* line 5 days after hatching. **B)** Brain coronal section with an eGFP immunohistochemistry of kif5a:eGFP-sponge-29 F1 20 weeks-old *N. furzeri*. **C)** Western blot of IRP2 and *TFR1A* in 12 weeks old kif5a:sponge-29 (n=4) and wild-type (n=4) fish brain extracts. **D)** Representative Western blot of IRP2 in brain extracts of kif5a:sponge-29 at different ages (5, 12, 20, 27). In C-D a-TUBULIN was used as loading control. **E)** Densitometric analysis of IRP2 expression. The solid blue line represents expression in kif5a:eGFP-sponge-29 animals, values were normalized to the mean of 5 weeks values, n=3 biological replicates were used for each time point. ANOVA for trend n.s. The gray line represents wild type fish. **F)** Expression levels of iron management genes measured through RT-qPCR in kif5a:eGFP-sponge-29 as compared to wild type. *TFR1A* alone was found deregulated (*, P<0,05, Mann-Whitney’ s U-test). **G)** Brain non-heme iron content (μg/g wet tissue) in kif5a:eGFP-sponge-29 (n=5) compared to wild type (n=6) at age 12 weeks (*, P<0,05, Mann-Whitney’s U-test).

### MiR-29 antagonism induces aging phenotypes in *N. furzeri* brain

To further elucidate the physiological consequences of miR-29 deficiency and, in turn, accelerated iron accumulation in adult brain, we analyzed the global gene expression profile through RNA-seq. We compared the expression profile of 12 weeks old transgenic fish with 12 weeks old wild type fish, using four biological replicates for experimental group. We detected 1307 differentially-expressed genes (DEGs, FDR < 0.05, edgeR). We performed a KEGG pathways analysis (FDR<0,05) on all DEGs. Among up-regulated DEGs we found an over-representation of genes involved in energy production processes such as oxidative phosphorylation and TCA cycle, including genes that encode for proteins known to be influenced by iron. In particular, Oexle et al. (1999) showed that iron induces aconitase 2 (*ACO2*), isocitrate dehydrogenase 3 (NAD+) beta (*IDH3B*), succinate dehydrogenase complex, subunit A (*SDHA*), all these four genes are up-regulated in transgenic fish (fig. 6A) (see also KEGG pathways graphical maps, appendix S1). Moreover, among up-regulated DEGs, we found also ribosome biogenesis, lysosome, phagosome and endoplasmic reticulum functions as well as downstream p53 effectors (fig. 6A) and an overrepresentation among down-regulated DEGs for genes involved in intracellular and extracellular remodeling, circadian rhythm and the following pathways: JAK-STAT, MAPK, WNT and Notch (fig. 6B; appendix S1). Several of these pathways were previously reported to be regulated in the *N. furzeri* brain during aging (see Baumgart M et al., 2014; Reichwald K et al., 2015). Remarkably, the majority of these showed the same direction of regulation during aging and in response to miR-29 antagonism (arrowed asterisk, fig. 6A-B) and only a minority, showed opposite directions of regulation (black asterisk, fig. 6A-B). Moreover, in a longitudinal study of *N. furzeri*, Baumgart et al. (2016) reported that higher expression of genes belonging to the oxidative phosphorylation, phagosome, lysosome, ribosome biogenesis and RNA transport pathways is negatively correlated with lifespan. Instead, higher expression of ECM-receptor interaction genes is positively correlated with lifespan (Baumgart M et al., 2016). We found that miR-29 antagonism induces an up-regulation of those pathways negatively correlated with lifespan (marked with a grey minus in fig. 6A-B), with the exception of RNA transport and spliceosome pathways and a down-regulation of the only pathway positively correlated with lifespan (marked with a plus in fig. 6A-B). Among up regulated DEGs, we found 25 and genes positively correlated and 3 negatively correlated with lifespan (P=0,0007, Fisher exact test, fig. 6C), among down-regulated DEGs, we found 9 genes positively correlated and 15 negatively correlated with lifespan (P=0,0049, Fisher exact test, fig. 6C). We interpreted these data as a signature of accelerated aging. To corroborate our hypothesis, we repeated the analysis at the gene level. We intersected the sets of DEGs of sponge-29 animals and aging (obtained by comparing gene expression of 5 weeks vs. 39 weeks old fish; Baumgart et al., 2014; Reichwald et al., 2015). The intersection contains 525 genes (fig. 6D) and 456 (~87%) show the same direction of regulation in aging and miR-29 sponge (p<10^-16^, *χ*^2^ test). Of those, 225 are up-regulated (Type 1 genes, fig. 6E) and 231 are down-regulated (type 3 genes, fig. 6E) in both cases. Only 68 genes (~13%) showed opposite regulation: 34 down-regulated in sponge-29 and up-regulated during aging (type 2, fig. 6E) and 34 with the opposite behavior (type 4, fig. 6E). Surprisingly, type 4 genes contained, in addition to *TFR1A*, other iron management genes such us the ferroxidase ceruloplasmin (CP) and transferrin a (TFA), confirming again the relationship between miR-29 and iron (the complete gene list is reported in supplementary table 1). Furthermore, we assessed the expression profile of those genes with a conserved predicted binding site for miR-29. To this end, we retrieved a list of D. rerio 548 predicted targets (score ≤ -0,30) from TargetScanFish (Ulitsky et al., 2012). Out of those, 38 genes were DEGs in kif5a:sponge-29 *N. furzeri* fish: 28 up-regulated and 10 down-regulated (fig. S6A). 22/28 *N. furzeri* ortholog up-regulated genes exhibited a conserved binding site (table fig. S6A). For 10/28, the miR-29 predicted binding site was conserved also in the mouse and human orthologs and for 5 of these an interaction with miR-29 was observed also by cross-linking immunoprecipitation (CLIP)-seq (fig. S6A). Here, as expected in accordance with the literature, we found genes involved in epigenetic reprogramming: Lysine-Specific Demethylase 6B (*KDM6BB*), DOT1-like, histone H3 methyltransferase (*DOT1L*), Enhancer of polycomb homolog 1 (*EPC1*), glutamate metabolism: Glutamate receptor-interacting protein 1 (*GRIP1*), IGF1 signaling: insulin-like growth factor binding protein 2a (*IGFBP2A*). Curiously, among down-regulated DEGs we found well-known mammalian miR-29 targets like: Ten-eleven translocation methylcytosine dioxygenase 3 (*TET3*), DNA (cytosine-5)-methyltransferase 3ab (*DNMT3AB*) and collagen type IV alpha 1 (*COL4A1*). Thus, we examined which was the effect of aging on conserved miR-29 predicted targets. Of 22 up-regulated targets 6/22 were found in type 4 dataset, 12/22 were not regulated by aging, indicating that for those miR-29 could directly influences their expression at transcriptional level. On the other hand 4/22 up-regulated and 9/10 of down-regulated targets were found respectively in type 1 and type 3 dataset (fig. S6A), indicating that probably the global effect of miR-29 depletion on aging is prevalent over the target-specific effect of miR-29. Finally, we analyzed 12 weeks old brains of kif5a:sponge-29 and wild type fish at histological level. Kif5a:sponge-29 fish exhibited an increased accumulation of the aging marker lipofuscin as compared to wild-type (P<0,01, fig. 7A-B), a marker of oxidative damage especially against cell or mitochondria membranes (Brunk and Terman, 2002) that is up-regulated in conditions of iron overload (Johnston and Milward, 2010; Johnston et al., 2012). Moreover, we observed an increased immunoreactivity for glial markers GFAP and S100β (P<0,05, fig. 7C-D), this data was also supported by RT-qPCR analysis and RNA-seq that both revealed an increased expression for relative mRNA genes in kif5a:sponge-29 fish (P<0,05, fig. S5B). Overall, these data strongly indicated that miR-29 family regulates brain iron metabolism during adult life and its deficiency induces an acceleration of some aspects of the aging phenotype.

**Figure 6.**
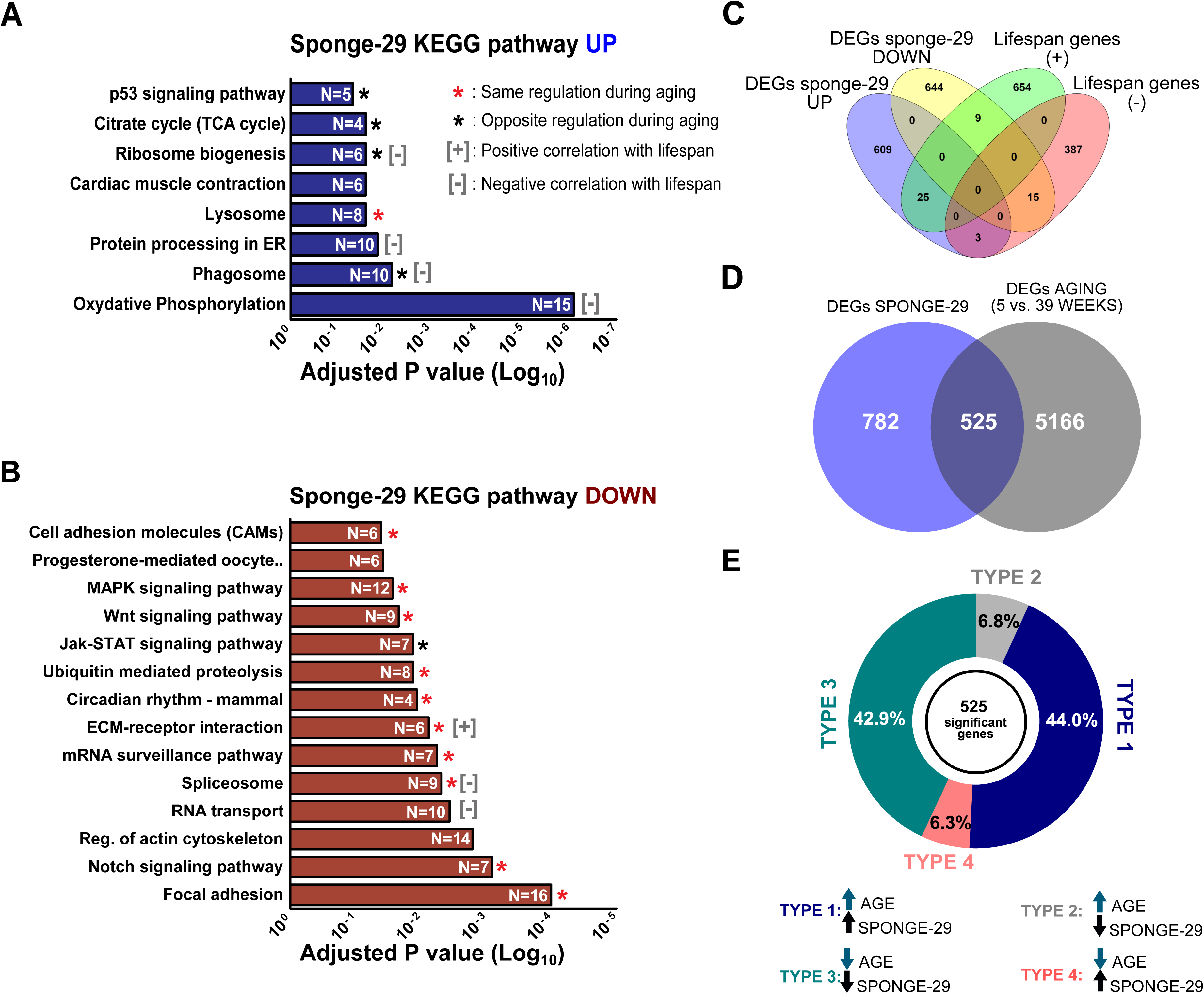
<u>MiR-29 antagonism accelerates aging rate</u> **A-B)** KEGG pathway overrepresentation analysis on DEGs (FDR≤0,01). The red and black asterisks indicate categories of DEGs regulated with age in the brain (from Reichwald et al., 2015). Red colored asterisk indicates categories that exhibit the same direction of regulation both in aging and sponge-29. Black colored asterisk indicates those with opposite regulation. Gray plus and minus symbols represent categories that positively and negatively correlates with lifespan respectively (from Baumgart et al., 2016) **C)** Venn diagram illustrating the intersection between up and downregulated DEGs in sponge-29 and genes positively and negatively correlated with lifespan respectively (P=0,007 and P=0,0049, Fisher exact test) **D)** Venn diagram illustrating the intersection between DEGs during aging and DEGs in sponge-29. **E)** Correlation analysis of fold-changes of genes in the intersection shown in (D). Pie chart represents the fraction of genes coherently or incoherently regulated by aging and sponge-29.

**Figure 7.**
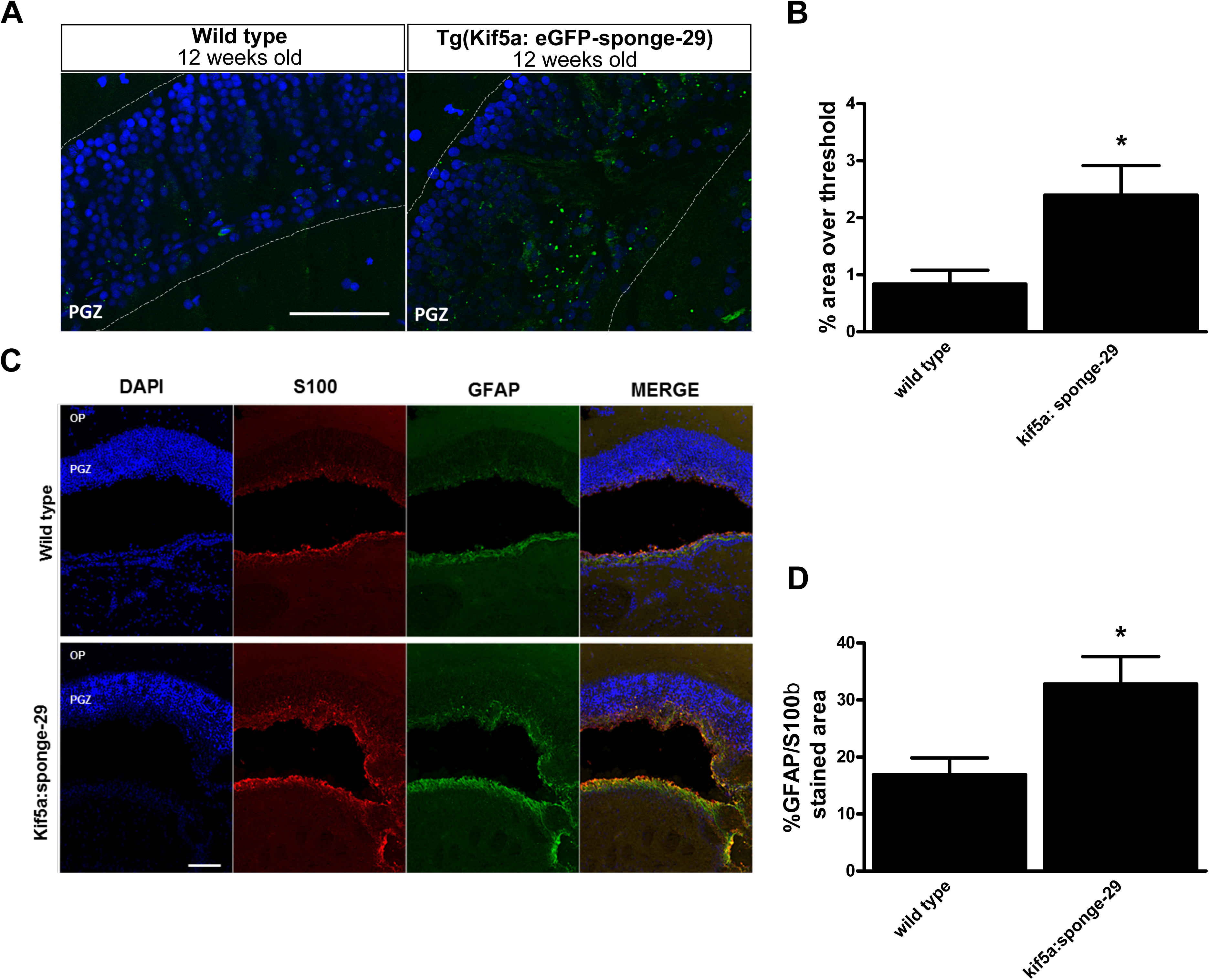
<u>miR-29 deficiency induces lipofuscin accumulation and gliosis</u> **A)** Representative images of lipofuscin accumulation in the optic tectum of 12 weeks old kif5a:eGFP-sponge-29 and wild type fish brains. Lipofuscin auto-fluorescent granules (green) were detected with ApoTome microscope, counterstained with DAPI (blue). Scale bar: 20μm. **B)** Quantification of lipofuscin density based on percentage of area over threshold, n=6 (*, P<0,05; Mann-Whitney’ s U-test). **C)** Representative images of immunoreactivity for GFAP and S100β markers in the optic tectum of 12 weeks old kif5a:eGFP-sponge-29 and wild type fish brains. **D)** Quantification of GFAP/S100β intensity per area of section. The analysis was performed in n=5 wild type and n=4 kif5a:sponge-29 fish brains (*, P<0,05, Mann-Whitney’ s U-test). Scale bar 100μm.

## Discussion

In this study, we investigated the effects of miR-29 regulation on iron metabolism during adult life in the brain of the short-lived fish *N. furzeri*. Expression of miR-29 is known to be upregulated during brain aging, here we report that: i) brain iron content increases with age; ii) miR-29 expression is induced by iron overload; iii) miR-29 negatively regulates the expression of *IREB2* gene in neurons during adult life; iv) miR-29 antagonism results in a chronic up-regulation of IRP2 and *TFR1A* resulting in increased brain iron deposition; v) transcriptome profile revealed an increased expression of TCA cycle and oxidative phosphorylation genes along with enhanced aging-related phenotype upon miR-29 antagonism vi) histological analysis revealed enhanced gliosis and lipofuscin deposition. These data demonstrate that up-regulation of miR-29 during aging is a compensatory response to limit an excessive age-dependent iron accumulation which can actively contribute to cell and tissues decline

IRP2 is an RNA-binding protein that acts as master regulator of intracellular iron availability in the brain, therefore influencing both energy production and redox status of cells. IRP2 is rapidly degraded when iron increases via allosteric activation of FBXL5 and this regulation defines a set point for physiological intracellular iron concentrations. Despite this rapid negative feedback regulation, iron accumulates in the brain during aging in *N*. *furzeri* as previously shown in mammals and it likely contributes to aging-induced dysfunctions. We also showed that, in fish, IRP2 protein, but not *IREB2* transcript, undergoes an age-dependent down-regulation but not its negative regulator FBXL5. Further, miR-29 directly targets *IREB2* in fish and mouse, it is up-regulated during aging and IRP2 is in turn up-regulated upon miR-29 antagonism, suggesting that under physiological conditions miR-29 exerts a translational control over IRP2 expression. This control, that requires transcriptional regulation of miR-29, obviously operates on a different time scale with respect to FBXL5-mediated degradation. Indeed, the increase of miR-29 upon iron overload has a kinetic of days, while iron accumulates within hours. Moreover, miR-29 expression level seems affected by iron overload, but not by iron depletion. In fact, reducing iron by 25% with DFO does not change miR-29 expression in the following 72 hours. The same dose of DFO, however, reduced miR-29 up-regulation if provided hours after iron injections when intracellular iron levels are already within the pathological range. Finally miR-29 antagonism chronically enhances cellular iron intake. Although the FBXL5 mechanism is present in the aged brain, it is insufficient to contrast a persistent increased iron level that in turn can exacerbate damage accumulation rate accelerating the aging process. Therefore, we suggest that increased miR-29 expression is a compensatory response to reduce iron intake when iron deposits as a consequence of age-related changes of cellular physiology.

Here, for the first time, we report the case of vertebrate system where neuronal specific perturbation of a microRNA induced iron imbalance and accelerated expression of aging phenotypes, both at the level of global gene expression and histological markers. Although we do not provide a formal proof for a causal relationship between increased iron and accelerated aging, this hypothesis is strongly supported by experiments in *C. elegans*. Klang et al. (2014) indeed reported that in *C. elegans* dietary iron significantly accelerates the aging-related phenotype, reduces the lifespan expectancy and increases age-dependent protein aggregation. We cannot exclude, however, that the effects of miR-29 antagonism on aging are not entirely mediated by iron and also involve other pathways that are directly targeted by this pleiotropic miRNA (e.g: epigenetic reprogramming, matrix remodeling). Overall, these data suggest that: i) loss of iron homeostasis gives a significant contribution to the normal aging process in vertebrates. ii) miR-29 is an important hub for preventing aging effects and iii) iron metabolism regulation is part of the protective action exerted by miR-29. Our data are in agreement with several independent reports on the protective role of this microRNA, for instance miR-29 promotes neuronal survival in conditions of acute neuronal injury such as ischemia (Khanna et al., 2013), miR-29 deficiency in mice induces neuronal loss in the cerebellum (Papadopoulou et al., 2015), an area where miR-29 is strongly expressed in *N. furzeri* as well, and most importantly excessive iron accumulation and down-regulation of miR-29 are both associated to Alzheimer’s disease (Smith et al., 1997; Hebert et al., 2008; Smith et al., 2010; Wang et al., 2011). Our data suggest that these two phenomena are linked and iron accumulation in AD is a consequence of reduced miR-29. All these data combined with the evolutionary conserved age-dependent regulation of miR-29 reinforce the general idea that it exerts a protective action in the aging brain.

Mitochondria are the main iron users and play a pivotal role in the aging process, although we not investigated mitochondria physiology in miR-29 downregulated animals, here we show that TCA cycle and oxidative phosphorylation genes are over-represented among up regulated genes upon miR-29 deficiency. Strikingly, insoluble proteins upon iron overload, as reported by Klang et al., (2014), are also enriched for TCA cycle and oxidative phosphorylation enzymes, suggesting that this transcriptional regulation is a compensatory response to iron-induced damages on mitochondrial proteins. These data suggest that miR-29 indirectly promotes mitochondrial function.

### Conclusions

Age-related damage accumulation is an inescapable condition that tends to change cellular homeostasis, on the other hand cells tend to maintain their homeostasis inducing a progressive and adaptive response in order to counteract this inevitable process and preserve their physiological functions. Age-dependent up-regulation of miR-29 is part of this adaptive response, its deficiency leads to exacerbation of aging-induced damage (fig. 8) partly due to impaired iron homeostasis.

**Figure 8.**
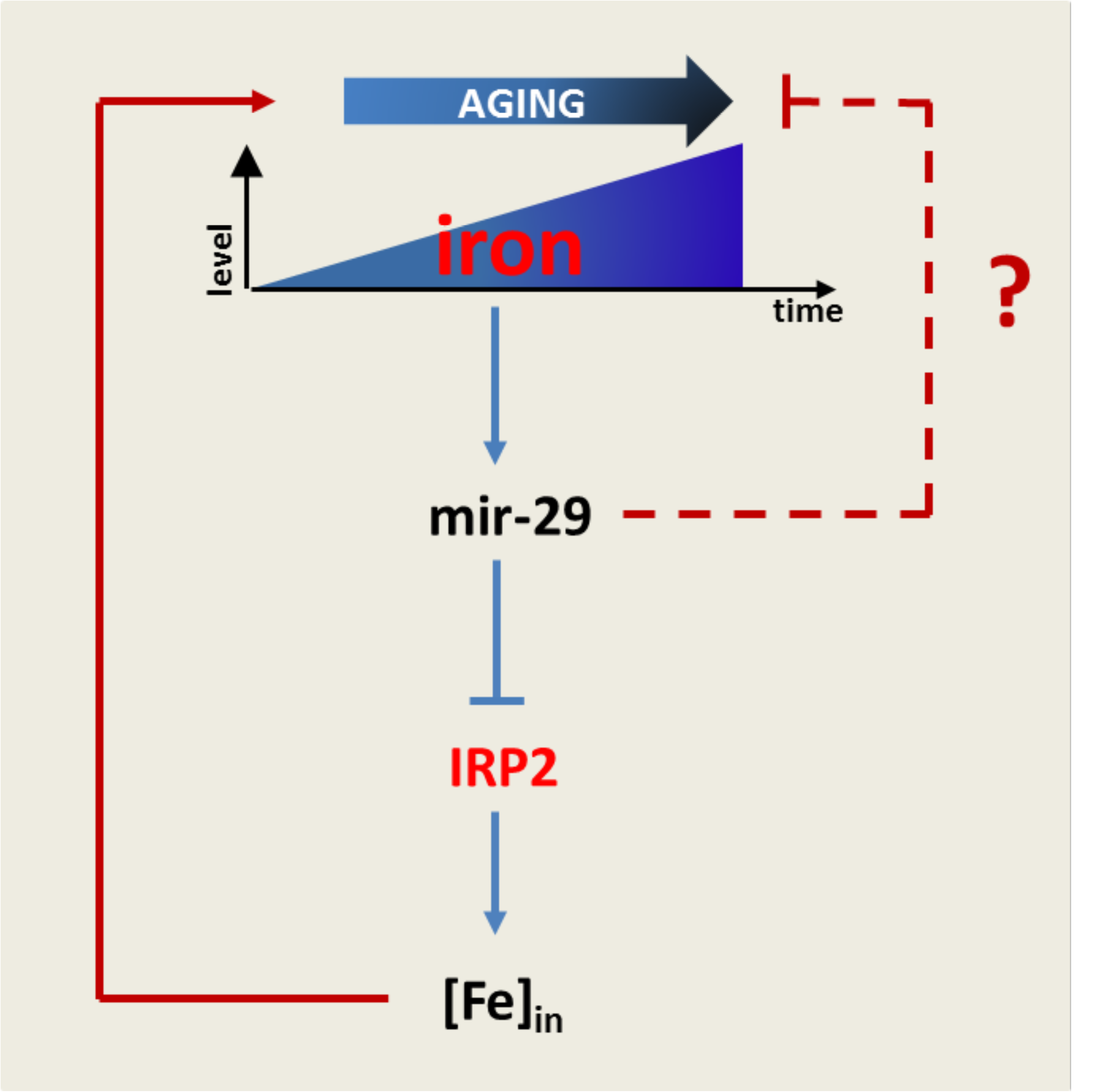
<u>miR-29 prevents early-onset aging process</u> A model of miR-29 function in iron metabolism. In response to the age-dependent iron accumulation, miR-29 expression increases and represses IRP2 by limiting excessive intracellular iron accumulation. This represents a compensatory response primed by cells in order to maintain cellular iron homeostasis and slow down the decline progression. Iron homeostasis regulation is likely one of several mechanisms by miR-29 prevents aging-induced physiological decline.

## Materials and methods

### Fish maintenance

*Nothobranchius furzeri:* all experiments were performed on group-house *N. furzeri* of the MZM-04/10 strain. The fish used were raised in 35L tanks at 25°C and were fed two to three times a day with frozen Chironomus larvae or living nauplii of *Artemia salina*, depending on size. Eggs were collected by sieving the sand with a plastic net and kept in wet peat moss during developmental processes and diapauses. Embryos were hatched by flushing the peat with tap water at 16-18°C. Embryos were scooped with a cut plastic pipette and transferred to a clean vessel. Fry were fed with newly hatched Artemia *nauplii* for the first 2 weeks and then weaned with finely chopped Chironomus larvae.

*Danio Rerio:* all experiments were performed on the Ab strain. Fish were maintained at 28°C under continuous flow in zebrafish facility with automatic control for a 14-hour light and 10-hour dark cycle (Zebtech system). To generate embryos for injection, male and female fish were placed the night before injection in a one liter fish tank with the inner mesh and divider. Zebrafish embryos were obtained from natural spawning by removing the divider and light stimulation.

### Vectors design and generation of transgenic lines

For the generation of all transgenesis vectors, we used multisite gateway technology (Kwan MK et al., 2007). Design of vector (kif5aa: EGFPCAAX-SPONGE-29-SV40pa): mir-29 sponge consisted of 7 repetition of mir-29 complementary sequences with a mismatch of 4 nucleotides immediately after the seed sequence. The sponge sequence was previously chemically synthetized by Eurofin (Milan) then cloned in the 3′ entry p3E-polyA using the restriction site BamHI. 3,094 kb of zebrafish kif5aa promoter region (Source: ZFIN;Acc:ZDB-GENE-070912-141) was amplified by PCR from genomic DNA, (from -2969 to +125 bp including a small portion of the first exon, before the ATG), using primers Kif5a-Sall f: (5′-gtcgac - GTTGTCCAGCACGGATGTATAGGTA-3′) and Kif5a-XmaI r:(5′-cccggg-ATAGCGGGGCAGAGGACGGCAG-3′) and cloned into gateway 5′-entry p5E-MCS. Then, the multisite gateway recombination reaction was performed as described in the Invitrogen Multi-Site Gateway Manual, an equimolar amount of entry vectors, 20 fg of each, (p5E-kif5aa, pEM-EGFPCAAX, and p3E-sponge-20-SV40pa) and destination vector (pDESTol2CG2) were combined with LR Clonase II Plus enzyme mix. For the generation of (actb2: EGFPCAAX-SPONGE-29-SV40pa) was used as 5′entry p5-bactin2 (from Tol2kit v1.2) together with the other plasmids following the protocol mentioned above. All these vectors also contain the eGFP reporter gene driven by the zebrafish cardiac myosin light chain (cmlc2) promoter. All vectors generated were extracted and purified using Qiagen plasmid midi kit (toxin free). Eggs were harvested and injected at zygote stage as described in the following protocols (Valenzano DR *et al*., 2011 and Xu Q, 1999 for *N. furzeri* and *D. rerio* microinjections respectively). For microinjection borosilicate microcapillaries were used. Capillaries were pulled with a micropipette puller (P97, Sutter Instrument). The needles were filled with 2μl of water solution containing 20-30 ng/μl of plasmid DNA, 20-30 ng/μl of Tol2 transposase mRNA, 0,4M KCl and 1% phenol red as visual control of successful injections. The embryos were injected with ~2nL of mix solution and the drop volume was estimated under a microscope using a calibrated slide.). Injections were performed under the Nikon C-PS stereoscope. F0 fishes were screened under fluorescence microscope at 3-4^th^ day after hatching (*N. furzeri*) and at 24hpf for zebrafish larvae. Three different founders were selected for line establishment.

### *IREB2* 3′UTR vectors assembly and mir-29 targeting validation

In order to experimentally validate *IREB2* as direct mir-29 target in fishes, 671bp and 633 bp from *IREB2* 3′UTR of *N. furzeri* and *D. rerio*, respectively, were amplified by PCR from cDNA of both species (*N. furzeri* reference: Nofu_GRZ_cDNA_3_0193494, *D. rerio* reference: ZFIN; Acc:ZDB-GENE-051205-1) using the primers nf*IREB2*-3′UTR-BamH-F: (5′-ttcgggggatccCATGTTTGACTCTGAGAAGGAC-3′) and nf*IREB2*-3′UTR-BamH-R:(5′-ttcgggggatccGTTCTGTGCCCAGTTTGCCC-3′) for *N. furzeri* and Dr*IRBB2*-3′UTR-BamH-F: (5′-gtcacgC-GGATCCTCACATGGACCTCTGAACACC-3′) and Dr*IRBB2*-3′UTR-BamH-R: (5′-gatcggc-GGATCCCACAGCGAAAGTATCACAGCC-3′) for *D. rerio* and cloned in the 3′ entry p3E-polyA, then p5E-CMV/SP6, pME-EGFPCAAX, p3E-IREB2-3′UTR-polyA and pDESTol2CG2 were combined using multisite gateway (see above). Finally, the plasmid producing fluorescence standard control was assembled combining p5E-CMV/SP6, pME-RFP, p3E-polyA and pDESTol2CG2. We renamed these plasmids as N.f CMV/SP6: eGFP-IREB2-pA, D.r CMV/SP6: eGFP-IREB2-pA and CMV/SP6: RFP-pA. For all of these reporters, *in vitro* transcription was carried out as described below. About 2nl of solution containing 100μg of egfp- *IREB2* sensor mRNA, 100μg of RFP standard, 100μg of Dre-mir-29a mimic (Qiagen) and 100μg of Dre-mir-29b mimic (exiqon) was microinjected in the zygote stage embryos (for the control embryos was assembled the same mix without microRNA mimic). At 24hpf, 40-50 embryos were collected from each experimental condition, dechorionated and dissociated with trypsin 10X at 37°C 15 min. Dissociated cell were diluted in PBS 1x, loaded in the flow cytometer machine (FacsCalibur, Becton-Dickinson) and the eGFP/RFP fluorescence ratio was analyzed and quantified. For *mmu-IREB2* 3′UTR validation we used dual luciferase reporter assay system (Promega), a sequence of *mmu-IREB2* 3′utr (transcript ID: ENSMUST00000034843) containing the putative mir-29 binding site was cloned in pMIR-report using the restriction enzymes Spel and Mlul. Mmu-mir-29a/b1 precursor was cloned in CMV/SP6: RFP-Pa vector, downstream of RFP using the restriction enzyme MluI. Transfection on Hek293 was performed using lipofectamine 2000 (Invitrogen). CMV:RFP-mir-29b/c precursor-polyA-tail was co-transfected with pMIR-report and Renilla control vector following the manufacturer’s instructions. In the control experiment was co-trasfected the same vector without mir-29a/b1 precursor sequence. 24 hours after transfection both Renilla and Firefly luciferase activity were measured using the luminometer (GloMax 96 microplate Luminometer w/dual injectors). Mutant vectors were generated using the QuikChange II XL Site-Directed Mutagenesis kit (Stratagene).

### In vitro RNA and DIG-labeled probes synthesis

The Tol2 mRNA was transcribed from the pCS-TP plasmid whereas eGFP-IREB2-3′utr and the standard fluorescence control mRNAs were transcribed from plasmids: N.f CMV/SP6: eGFP-*IREB2-pA*, D.r CMV/SP6: eGFP-IREB2-pA and CMV/SP6: RFP-pA respectively. All were previously linearized using NotI and purified with Wizard^®^ SV Gel and PCR Clean-Up System (Promega). Then 1μg of each linearized plasmid was transcribed using the mMESSAGE mMACHINE SP6 kit (Ambion) according to the manifacturer’s protocol. For DIG-labeled probe synthesis, initially the probes were amplified by PCR from cDNA using a reverse primer carrying a T7 promoter sequence in its 5′end. 200ng of PCR product, previously purified with Wizard^®^ SV Gel and PCR Clean-Up System (Promega), were directly transcribed using T7 enzyme (Fermentas) and digoxygenated RNTP mix (Roche) 2 hours at 37°C. In vitro transcription mix were precipitated with 1/10 of volume of LiCl (5M) and 2,5 volumes of isopropanol, than washed with 70% ethanol and finally resuspended in nuclease-free water.

### Total RNA extraction and RT-qPCR

Dissected tissues were immediately put in 500 μl of Qiazol lysis reagent and manually homogenized with pounder. Total RNA was extracted using miRneasy mini kit (Qiagen) according to the manufacture’s protocol. 200ng of each RNA extraction was retrotranscribed for cDNA synthesis using mirscript II RT kit (Qiagen). qPCR was performed using Rotorgene 6000 (Corbet). PCR mix solutions were prepared using SsoAdvanced™ Universal SYBR^®^ Green Supermix (Biorad) and 4ng of cDNA for each sample as template. The relative gene quantification was calculated using the ΔΔCt method, as reference genes were used *TBP, ACTB2* and *GAPDH* for *N*.*furzeri, D. rerio* and mouse respectively.

### Histology and histochemistry

All the immunohistochemical procedures were performed on frozen tissue sections. Animals were killed with overdose of MS-222, brains were dissected and fixed in PFA 4%, washed in PBS 1X twice then equilibrated in sucrose 30% and embedded in Tissue-Tek OCT (Leica). Frozen tissues were cut with cryostat (Leica), 12-14 μm thick sections were immediately put on superfrost plus slides (Thermo Scientific), dried in the oven at 55°C for 1 hour. Sections were rehydrated in PBS 1x permeabilized with tritonX 0,3% and blocked in BSA 5%, goat serum 1%. Primary antibodies were all incubated over night at 4°C according to the following dilutions: Anti-IRP2 (1:500, Abcam), Anti-*TFRC* (1:500, Abcam), Anti-HuC/D (1:50, Invitrogen), Anti-eGFP (1:1000, Abcam), Anti-GFAP (1:400, Invitrogen), Anti-S100 (1:400, Dako). Secondary antibodies coupled to Alexa Fluor dye (488, 546, 635) were incubated 2 hours RT (1:500).

For *in situ* hybridizations, slides were incubated with proteinase K for 10 minutes at room temperature (RT)(1:80000; Fermentas 20mg/ml), post-fixed with PFA 4% 20 minutes RT, then were incubated with a digoxigenin (DIG(-labeled probes (60°C, ON). Immediately before incubation probes were put in hybridization solution denaturated at 98°C for 3 min. Sections were washed with SSC2x twice at 60°C for 15min. each, then with SSC 0,5x three times 10 min each RT and incubated with anti-Dig AP Fab (Roche; 1/2000 4°C ON) in blocking solution (Roche). The day after, slides were placed 20 min RT in TMN (Tris-MgCI2-NaCI buffer) with the addition of levamisole 1 mM (Sigma) in order to inhibit endogenous alkaline phosphatase and transferred in Fast red solution (Roche tablets). The staining was constantly monitored under epiflourescence microscope and blocked by washing in PBS 1X. Images were acquired using epifluorescence microscope (Nikon, Eclipse600) or confocal microscope (Leica DMIRE2).

For lipofuscin detection, unstained sections were deparaffinized and mounted using a water-based medium within DAPI (Invitrogen). An important property of lipofuscin is its broad autofluorescence. So it was acquired using Zeiss apotome(2) at an excitation wavelength of 488 and 550nm as well and under UV excitation was acquired DAPI staining. Images were analyzed using Image-j, threshold were determined by operator and applied to images to discriminate lipofuscin granules from background signal. The area occupied by granules was expressed as a percentage of total image area analyzed.

### Iron staining (Perl’s staining)

For iron staining, according with (Meguro R et al., 2007), with some adjustments, brains were dissected and fixed in PFA 4% and embedded in paraffin. 5-7 μm thick sections were deparaffinized and immersed in 4% ferrocyanide and 2% HCl for 1 hour at 37°C, then immersed in methanol containing 0,5 % H_2_O_2_, 0,01 NaN_3_ for 30 min RT, finally immersed in 0.1 M phosphate buffer containing DAB 0,05%, 0,005% H_2_O_2_ and 0,1% tritonX for 30 min. Sections were counterstained with hematoxylin, dehydrated and mounted.

### Non heme iron quantification

For iron quantification was followed the protocol published by (Rebouche JC. *et al*. 2003). Extracted tissues were put in a 1.5ml tube previously weighed and were burden again in order to precisely determine wet tissue weight. Homogenates were prepared in high-purity water (max resuspension volume 100μl). 50 μl of homogenate tissues were combined in a new 1.5ml tube with an equal volume of protein precipitation solution (1N HCl, 10% trichloroacetic acid) and placed in thermoblock at 95°C for 1 hour. Tubes were cooled RT for 10 min then were centrifuged for 15 min at 4°C. 70 μl of supernatant was collected from each tube sample and combined with an equal volume of chromogenic solution (Ferrozine 0,5 mM, ammonium acetate 1,5 M and Tioglicolic acid 0,1%). After 30 min absorbance was measured at 562 nm using spectrophotometer. Standard curve were prepared using 0, 0,5, 1, 2, 4, 8, 10, 20 μg/ml of iron standard solution (Sigma).

### Iron and drugs delivery

Fish were previously anesthetized with Trichaine methanesulfonate and weighed. A single dose respectively of 350 μg/g body weight iron dextran (Sigma, 100mg/ml), 30 μg/g body weight deferoxamine (DFO, Novartis) and 50 μ/g body weight 4-OH-Tempol (Sigma) was injected intraperitonealy. Injections were performed under a stereo microscope (Leica) using Amilton syringes 10μl, control animals were sham-injected with saline solution. At 4, 8, 12, 24, 48 and 72 hours after injection brains were removed from anesthetized animals and put in a previously weighed tube for iron measurement or immediately frozen in liquid nitrogen for the following RNA extraction.

### Western blot

Samples were lysed in RIPA-buffer (50 mM Tris-HCl, pH 7.5, 150 mM NaCl, 1% Triton X-100, 0.1% SDS, 0,5% deoxycolic acid) containing protease inhibitor (Complete Protease Inhibitor Cocktail Tablets, Roche Diagnostics) and centrifuged at 12300 rpm for 10 min at 4°C and supernatants were collected. Total protein concentration was determined by BCA protein assay (Pierce). Aliquots of homogenate with equal protein concentrations were separated in 10% acrylamide gel and transferred to nitrocellulose membranes by mini trans-blot (Bio-Rad). The membranes were blocked with milk (5% w/v) and probed with appropriate primary and secondary IgG-HRP conjugated antibodies (Millipore). Enhanced chemiluminiscence detection system (GE Healthcare) was used for developing on autoradiography-films (GE Healthcare). Densitometric quantification was performed using image-j and normalized to the relative amount of (μIII-Tubulin and expressed as n-fold of control samples. In this study were used the following antibodies anti-IRP2 (Abcam 1:1000), anti-*TFRC* (Abcam 1:500).

### Cell culture and iron exposure

Following a previously published protocol (Bertacchi et al., Cell Mol Life Sci, 2013), murine ES cell line E14Tg2A was differentiated into cortical neurons. In brief, mouse embryonic stem cells (mESC) were cultured in a chemically defined medium on laminin-coated culture dishes for 20 days. mESC-derived neurons were then treated with iron dextran (50 ug/mL, 100 ug/mL and 200 ug/mL) and Iron(II) sulfate (25 uM, 50 uM and 100 uM) in Neurobasal Medium (Thermo Fisher Scientific) supplemented with B27 (Thermo Fisher Scientific) for 72 hrs, medium was changed daily. Subsequently, two wells of six-well plate were pooled for each thesis; RNA extraction and PCR analysis were carried out from three biological replicates.

### RNA-sequencing and analysis

RNA was extracted using Quaziol (Qiagen). Library preparation using Illumina’s TruSeq RNA sample prep kit v2 and sequenced on Illumina HiSeq2500, 50bp single-read mode in multiplexing obtaining around 50-40 mio reads per sample. Read mapping was performed using Tophat 2.0.6 (Kim D et al., 2013) and featureCounts v1.4.3-p1 (Liao Y et al., 2014) using as refences the *N. furzeri* genome (Reichwald K et al., 2015) or Zv9.73. Differential gene expression analysis was performed using R software, for DEGs identification was used the statistical test of edgeR package (Robinson et al., 2010). For multiple testing correction a false discovery rate <0.05 was chosen. KEGG pathway analysis was performed using the software Web-based gene set analysis tool kit (WebGestalt) (Zhang et al., 2005) using an FDR<0,05.

### Ethic authorization for animal experimentation

Procedure for animal husbandry and breeding were authorized by the Italian Ministery of Health (Aut N. 96/2003-A).

Animal experiment procedure were specifically approved by the local ethical committee and the Italian Ministery of Health (Aut. N. 1314/2015-PR).

## Acknowledgments

We thank Matthias Platzer for access to the sequencing facility and support through this project, Letizia Pitto for access to the zebrafish fishroom,Uwe Menzel for help in analysis, Mirko Mutalipassi for technical assistance in the fishroom, Marco Matteucci for access to the histology facility and Nicola Carucci, Giovanna Testa and Caterina Rizzi for sharing mouse samples.

This work was supported by internal grant of Scuola Normale Superiore and a grant of the Bundesamt für Bildung und Forschung (JenAge; BMBF, support codes: 0315581A).

## References

1. Abboud S, Haile DJ. (2000) A novel mammalian iron-regulated protein involved in intracellular iron metabolism. J Biol Chem. 275(26):19906–12.

2. Agarwal V., Bell G. W., Nam J. W., & Bartel D. P. (2015). Predicting effective microRNA target sites in mammalian mRNAs. eLife.

3. Asano T, Koike M, Sakata S, Takeda Y, Nakagawa T, Hatano T, Ohashi S, Funayama M, Yoshimi K, Asanuma M, Toyokuni S, Mochizuki H, Uchiyama Y, Hattori N, Iwai K. (2015) Possible involvement of iron-induced oxidative insults in neurodegeneration. Neurosci Lett. 588:29-35.

4. Bartzokis G., Beckson M., Hance D. B., Marx P., Foster J. A., Marder S. R. (1997). MR evaluation of age-related increase of brain iron in young adult and older normal males.

5. Baumgart M, Groth M, Priebe S, Appelt J, Guthke R, Platzer M, Cellerino A. (2012) Age-dependent regulation of tumor-related microRNAs in the brain of the annual fish Nothobranchius furzeri. Mech Ageing Dev. 133(5):226-33.

6. Baumgart M, Priebe S, Groth M, Hartmann N, Menzel U, Pandolfini L, Koch P, Felder M, Ristow M, Englert C, Guthke R, Platzer M, Cellerino A (2016) Longitudinal transcriptional analysis of vertebrate aging identifies mitochondrial 2 complex I as a small molecule-sensitive modifier of lifespan. Cell system (in press)

7. Baumgart M., Groth M., Priebe S., Savino A., Testa G., Dix A., et al Cellerino, A. (2014). RNA-seq of the aging brain in the short-lived fish N. furzeri - conserved pathways and novel genes associated with neurogenesis. Aging Cell, 13(6), 965–974.

8. Bertacchi M., Pandolfini L., Murenu E., Viegi A., Capsoni S., Cellerino A., et al Cremisi, F. (2013). The positional identity of mouse ES cell-generated neurons is affected by BMP signaling. Cellular and Molecular Life Sciences, 70(6), 1095–1111.

9. Betel D., Koppal A., Agius P., Sander C., & Leslie C. (2010). Comprehensive modeling of microRNA targets predicts functional non-conserved and non-canonical sites. Genome Biology, 11(8), R90.

10. Boon RA, Seeger T, Heydt S, Fischer A, Hergenreider E, Horrevoets AJ, Vinciguerra M, Rosenthal N, Sciacca S, Pilato M, van Heijningen P, Essers J, Brandes RP, Zeiher AM, Dimmeler S. (2011) Circ Res. MicroRNA-29 in aortic dilation: implications for aneurysm formation. 109(10):1115–9.

11. Boudreau RL, Jiang P, Gilmore BL, Spengler RM, Tirabassi R, Nelson JA, Ross CA, Xing Y, Davidson BL. (2014) Transcriptome-wide discovery of microRNA binding sites in human brain. Neuron. 81(2):294–305.

12. Brunk U. T., & Terman A. (2002). Lipofuscin: Mechanisms of age-related accumulation and influence on cell function. Free Radical Biology and Medicine. 5849(02)00959-0.

13. Cellerino A., Valenzano D. R., & Reichard M. (2015). From the bush to the bench: The annual Nothobranchius fishes as a new model system in biology. Biological Reviews.

14. Chen P. Y., Manninga H., Slanchev K., Chien M., Russo J. J., Ju J., et al Tuschl, T. (2005). The developmental miRNA profiles of zebrafish as determined by small RNA cloning. Genes and Development, 19(11), 1288–1293.

15. Crespo ÂC, Silva B, Marques L, Marcelino E, Maruta C, Costa S, Timóteo A, Vilares A, Couto FS, Faustino P, Correia AP, Verdelho A, Porto G, Guerreiro M, Herrero A, Costa C, de Mendonça A, Costa L, Martins M. (2014) Genetic and biochemical markers in patients with Alzheimer’s disease support a concerted systemic iron homeostasis dysregulation. Neurobiol Aging. 35(4):777–85.

16. Cushing L, Costinean S, Xu W, Jiang Z, Madden L, Kuang P, Huang J, Weisman A, Hata A, Croce CM, Lü J. (2015) Disruption of miR-29 Leads to Aberrant Differentiation of Smooth Muscle Cells Selectively Associated with Distal Lung Vasculature. PLoS Genet, 11(5), e1005238.

17. Di Cicco E, Tozzini ET, Rossi G, Cellerino A. (2011) The short-lived annual fish Nothobranchius furzeri shows a typical teleost aging process reinforced by high incidence of age-dependent neoplasias. Exp Gerontol. 46(4):249–56.

18. Dixon SJ, Stockwell BR. (2014) The role of iron and reactive oxygen species in cell death. Nat Chem Biol. 2014 Jan;10(1):9-17.

19. Dlouhy AC, Outten CE. The iron metallome in eukaryotic organisms. Metal Ions Life Sci. 2013;12:241-278.

20. Duce JA, Tsatsanis A, Cater MA, James SA, Robb E, Wikhe K, Leong SL, Perez K, Johanssen T, Greenough MA, Cho HH, Galatis D, Moir RD, Masters CL, McLean C, Tanzi RE, Cappai R, Barnham KJ, Ciccotosto GD, Rogers JT, Bush AI. (2010) Iron-export ferroxidase activity of β-amyloid precursor protein is inhibited by zinc in Alzheimer’s disease. Cell. 142(6):857–67.

21. Eichhorn S. W., Guo H., McGeary S. E., Rodriguez-Mias, R. A., Shin C., Baek, D., et al Bartel, D. P. (2014). MRNA Destabilization Is the dominant effect of mammalian microRNAs by the time substantial repression ensues. Molecular Cell, 56(1), 104–115.

22. Febbraro F, Giorgi M, Caldarola S, Loreni F, and Romero-Ramos, M (2012) a-Synuclein expression is modulated at the translational level by iron. Neuroreport 23, 576–580.

23. Fenn AM, Smith KM, Lovett-Racke AE, Guerau-de-Arellano M, Whitacre CC, Godbout JP. (2013) Increased micro-RNA 29b in the aged brain correlates with the reduction of insulin-like growth factor-1 and fractalkine ligand. Neurobiol Aging. 34(12):2748–58.

24. Friedlich AL, Tanzi RE, and Rogers JT (2007) The 5-untranslated region of Parkinson’s disease alpha- synuclein messengerRNA contains a predicted iron responsive element. Mol. Psychiatry 12, 222–223.

25. Garrick M. D., Dolan K. G., Horbinski C., Ghio A. J., Higgins D., Porubcin M., et al. (2003). DMT1: a mammalian transporter for multiple metals. Biometals 16, 41–54.

26. Gerhard G. S., Kauffman E. J., Wang X., Stewart R., Moore J. L., Kasales C. J., et al Cheng, K. C. (2002). Life spans and senescent phenotypes in two strains of Zebrafish (Danio rerio). Experimental Gerontology, 37 (8-9), 1055-1068.

27. Haile DJ, Rouault TA, C K Tang, J Chin, J B Harford, and R D Klausner (1992) Reciprocal control of RNA-binding and aconitase activity in the regulation of the iron-responsive element binding protein: role of the iron-sulfur cluster. Proc Natl Acad Sci USA. 89(16):7536–7540.

28. Hebert SS, Horre K, Nicolai L, Papadopoulou AS, Mandemakers W, Silahtaroglu AN, Kauppinen S, Delacourte A, De Strooper B. (2008) Loss of microRNA cluster miR-29a/b-1 in sporadic Alzheimer’s disease correlates with increased BACE1/beta-secretase expression. Proc Natl Acad Sci U S A. 105(17):6415–20.

29. Johnstone D., & Milward E. A. (2010). Genome-wide microarray analysis of brain gene expression in mice on a short-term high iron diet. Neurochemistry International, 56 (6-7), 856-863.

30. Johnstone D., Graham R. M., Trinder D., Delima R. D., Riveros C., Olynyk J. K., et al Milward, E. A. (2012). Brain transcriptome perturbations in the Hfe -/- mouse model of genetic iron loading. Brain Research, 1448, 144–152.

31. Khanna S, Rink C, Ghoorkhanian R, Gnyawali S, Heigel M, Wijesinghe DS, Chalfant CE, Chan YC, Banerjee i, Huang Y, Roy S, Sen CK. (2013) Loss of miR-29b following acute ischemic stroke contributes to neural cell death and infarct size. J Cereb Blood Flow Metab. 33(8):1197–206.

32. Kim D., Pertea G., Trapnell C., Pimentel H., Kelley R., and Salzberg, S.L. (2013). TopHat2: accurate alignment of transcriptomes in the presence of insertions, deletions and gene fusions. Genome Biol 14, R36.

33. Kiang I. M., Schilling B., Sorensen D. J., Sahu A. K., Kapahi P., Andersen J. K., et al Lithgow, G. J. (2014). Iron promotes protein insolubility and aging in C. elegans. Aging, 6(11), 975–991.

34. Kole AJ, Swahari V, Hammond SM, Deshmukh M. miR-29b is activated during neuronal maturation and targets BH3-only genes to restrict apoptosis. (2011) Genes Dev. 25(2):125-30.

35. Kwan KM, Fujimoto E, Grabher C, Mangum BD, Hardy ME, Campbell DS, Parant JM, Yost HJ, Kanki JP, Chien CB. (2007) The Tol2kit: a multisite gateway-based construction kit for Tol2 transposon transgenesis constructs. Dev Dyn. 236(11):3088-99.

36. Landgraf P, Rusu M, Sheridan R, Sewer A, Iovino N, Aravin A, Pfeffer S, Rice A, Kamphorst AO, Landthaler M, Lin C, Socci ND, Hermida L, Fulci V, Chiaretti S, Foà R, Schliwka J, Fuchs U, Novosel A, Müller RU, Schermer B, Bissels U, Inman J, Phan Q, Chien M, Weir DB, Choksi R, De Vita G, Frezzetti D, Trompeter HI, Hornung V, Teng G, Hartmann G, Palkovits M, Di Lauro R, Wernet P, Macino G, Rogler CE, Nagle JW, Ju J, Papavasiliou FN, Benzing T, Lichter P, Tam W, Brownstein MJ, Bosio A, Borkhardt A, Russo JJ, Sander C, Zavolan M, Tuschl T. (2007) A mammalian microRNA expression atlas based on small RNA library sequencing. Cell. 129(7):1401–14.

37. LaVaute T, Smith S, Cooperman S, Iwai K, Land W, Meyron-Holtz E, Drake SK, Miller G, Abu-Asab M, Tsokos M, Switzer R 3rd, Grinberg A, Love P, Tresser N, Rouault TA. (2001) Targeted deletion of the gene encoding iron regulatory protein-2 causes misregulation of iron metabolism and neurodegenerative disease in mice. Nat Genet. 27(2):209-14.

38. Li H., Mao S., Wang H., Zen K., Zhang, C., & Li L. (2014). MicroRNA-29a modulates axon branching by targeting doublecortin in primary neurons. Protein and Cell, 5(2), 160–169.

39. Li J. H., Liu S., Zhou H., Qu L. H., & Yang J. H. (2014). StarBase v2.0: Decoding miRNA-ceRNA, miRNA-ncRNA and protein-RNA interaction networks from large-scale CLIP-Seq data. Nucleic Acids Research, 42(D1).

40. Liao Y., Smyth G.K., and Shi, W. (2014). Feature Counts: an efficient general purpose program for assigning sequence reads to genomic features. Bioinformatics 30, 923–930.

41. Mantyh PW, Ghilardi JR, Rogers S, DeMaster E, Allen CJ, Stimson ER, Maggio JE (1993) Aluminum, iron, and zinc ions promote aggregation of physiological concentrations of beta-amyloid peptide. J Neurochem. 61:1171-1174.

42. McKie AT, Marciani P, Rolfs A, Brennan K, Wehr K, Barrow D, Miret S, Bomford A, Peters TJ, Farzaneh F, Hediger MA, Hentze MW, Simpson RJ. (2000) A novel duodenal iron-regulated transporter, IREG1, implicated in the basolateral transfer of iron to the circulation. Mol Cell. 5(2):299-309.

43. Meguro R, Asano Y, Odagiri S, Li C, Iwatsuki H, Shoumura K. (2007) Nonheme-iron histochemistry for light and electron microscopy: a historical, theoretical and technical review. Arch Histol Cytol. 70(1):1-19.

44. Meyron-Holtz EG1, Ghosh MC, Iwai K, LaVaute T, Brazzolotto X, Berger UV, Land W, Ollivierre-Wilson H, Grinberg A, Love P, Rouault TA. (2004) Genetic ablations of iron regulatory proteins 1 and 2 reveal why iron regulatory protein 2 dominates iron homeostasis. EMBO J. 23(2):386-95.

45. Oexle H, Gnaiger E, Weiss G. (1999) Iron-dependent changes in cellular energy metabolism: influence on citric acid cycle and oxidative phosphorylation. Biochim Biophys Acta. 1413(3):99–107.

46. Oshiro S, Morioka MS, Kikuchi M. (2011) Dysregulation of iron metabolism in Alzheimer’s disease, Parkinson’s disease, and amyotrophic lateral sclerosis. Adv Pharmacol Sci. 2011:378278.

47. Ouyang YB, Xu L, Lu Y, Sun X, Yue S, Xiong, Giffard RG. (2013) Astrocyte-enriched miR-29a targets PUMA and reduces neuronal vulnerability to forebrain ischemia. Glia. 61(11):1784-94.

48. Papadopoulou AS, Serneels L, Achsel T, Mandemakers W, Callaerts-Vegh Z, Dooley J, Lau P, Ayoubi T, Radaelli E, Spinazzi M, Neumann M, Hébert SS, Silahtaroglu A, Liston A, D’Hooge R, Glatzel M, De Strooper B. (2015) Deficiency of the miR-29a/b-1 cluster leads to ataxic features and cerebellar alterations in mice. Neurobiol Dis. 73:275-88.

49. Penke L, Valdés Hernández MC, Muñoz Maniega S, Gow AJ, Murray C, Starr JM, Bastin ME, Deary IJ, Wardlaw JM (2012) Brain iron deposits are associated with general cognitive ability and cognitive aging. Neurobiol Aging 33:510-517.

50. Podolska A, Kaczkowski B, Kamp Busk P, Søkilde R, Litman T, Fredholm M, Cirera S. (2011) MicroRNA expression profiling of the porcine developing brain. PLoS One. 26(1):e14494.

51. Rebouche CJ, Wilcox CL, Widness JA. (2003) Microanalysis of non-heme iron in animal tissues. J Biochem Biophys Methods. 58(3):239-51.

52. Reichwald K, Petzold A, Koch P, Downie BR, Hartmann N, Pietsch S, Baumgart M, Chalopin D, Felder M, Bens M, Sahm A, Szafranski K, Taudien S, Groth M1, Arisi I, Weise A, Bhatt SS, Sharma V, Kraus JM, Schmid F, Priebe S, Liehr T, Görlach M, Than ME, Hiller M, Kestler HA, Volff JN, Schartl M, Cellerino A, Englert C, Platzer M. (2015) Insights into Sex Chromosome Evolution and Aging from the Genome of a Short-Lived Fish. Cell. 16(6): 1527-38.

53. Robinson MD, McCarthy DJ and Smyth GK (2010). edgeR: a Bioconductor package for differential expression analysis of digital gene expression data. Bioinformatics. 26(1):139-40.

54. Roshan R., Shridhar S., Sarangdhar M. A., Banik A., Chawla M., Garg M., et al Pillai, B. (2014). Brain-specific knockdown of miR-29 results in neuronal cell death and ataxia in mice. RNA, 20(8), 1287–1297.

55. Rouault TA, Tong WH. (2005) Iron-sulphur cluster biogenesis and mitochondrial iron homeostasis. Nat Rev Mol Cell Biol. 6(4):4-345.

56. Rouault T. A. (2013). Iron metabolism in the CNS: implications for neurodegenerative diseases. Nature Reviews Neuroscience, 14(8), 551–564.

57. Salahudeen A. A., Thompson J. W., Ruiz J. C., Ma H. W., Kinch L. N., Li Q., Grishin N. V., Bruick R. K. (2009) An E3 ligase possessing an iron-responsive hemerythrin domain is a regulator of iron homeostasis. Science 326, 722-726.

58. Sanchez M1, Galy B, Schwanhaeusser B, Blake J, Bähr-Ivacevic T, Benes V, Selbach M, Muckenthaler MU, Hentze MW. (2011) Iron regulatory protein-1 and -2: transcriptome-wide definition of binding mRNAs and shaping of the cellular proteome by iron regulatory proteins. Blood. 24;118(22):e168-79.

59. Smith MA, Harris PL, Sayre LM, Perry G. (1997) Iron accumulation in Alzheimer disease is a source of redox-generated free radicals. Proc Natl Acad Sci USA. 94(18):9866–8.

60. Smith MA, Zhu X, Tabaton M, Liu G, McKeel DW Jr, Cohen ML, Wang X, Siedlak SL, Dwyer BE, Hayashi T, Nakamura M, Nunomura A, Perry G. (2010) Increased iron and free radical generation in preclinical Alzheimer disease and mild cognitive impairment. J Alzheimers Dis. 19(1):363-72.

61. Somel M, Guo S, Fu N, Yan Z, Hu HY, Xu Y, Yuan Y, Ning Z, Hu Y, Menzel C, Hu H, Lachmann M, Zeng R, Chen W, Khaitovich P. (2010) MicroRNA, mRNA, and protein expression link development and aging in human and macaque brain. Genome Res. 20,1207-1218.

62. Takahashi M, Eda A, Fukushima T, Hohjoh H (2012) Reduction of type IV collagen by upregulated miR-29 in normal elderly mouse and klotho-deficient, senescence-model mouse. PLoS One. 7(11):e48974.

63. Terzibasi E. T., Baumgart M., Battistoni, G., & Cellerino A. (2012). Adult neurogenesis in the short-lived teleost Nothobranchius furzeri: Localization of neurogenic niches, molecular characterization and effects of aging. Aging Cell, 11(2), 241–251.

64. Terzibasi E., Lefranois, C., Domenici P., Hartmann N., Graf M., & Cellerino A. (2009). Effects of dietary restriction on mortality and age-related phenotypes in the short-lived fish Nothobranchius furzeri. Aging Cell, 8(2), 88–99.

65. Terzibasi E., Valenzano D. R., Benedetti M., Roncaglia P., Cattaneo A., Domenici, L., & Cellerino A. (2008). Large differences in aging phenotype between strains of the short-lived annual fish Nothobranchius furzeri. PLoS ONE, 3(12).

66. Theil EC. (1990) Regulation of ferritin and transferrin receptor mRNAs. J Biol Chem. 265(9):4771–4.

67. Ugalde A. P., Ramsay A. J., de la Rosa, J., Varela I., Mariño G., Cadiñanos J., et al López-Otín, C. (2011). Aging and chronic DNA damage response activate a regulatory pathway involving miR-29 and p53. The EMBO Journal, 30(11), 2219–2232.

68. Ulitsky I., Shkumatava A., Jan C. H., Subtelny A. O., Koppstein D., Bell G. W., et al Bartel, D. P. (2012). Extensive alternative polyadenylation during zebrafish development. Genome Research, 22(10), 2054–2066.

69. Valenzano DR, Benayoun BA, Singh PP, Zhang E, Etter PD, Hu CK, Clément-Ziza M, Willemsen D, Cui R, Harel I, Machado BE, Yee MC, Sharp SC, Bustamante CD, Beyer A, Johnson EA, Brunet A. (2015) The African Turquoise Killifish Genome Provides Insights into Evolution and Genetic Architecture of Lifespan. Cell. 163(6):6-1539.

70. Valenzano DR, Sharp S, Brunet A. (2011) Transposon-Mediated Transgenesis in the Short-Lived African Killifish Nothobranchius furzeri, a Vertebrate Model for Aging. G3 (Bethesda). 1(7):531-8.

71. Valenzano DR, Terzibasi E, Cattaneo A, Domenici L, Cellerino A. (2006a) Temperature affects longevity and age-related locomotor and cognitive decay in the short-lived fish Nothobranchius furzeri. Aging Cell. 5(3):275-8.

72. Valenzano DR, Terzibasi E, Genade T, Cattaneo A, Domenici L, Cellerino A. (2006b) Resveratrol prolongs lifespan and retards the onset of age-related markers in a short-lived vertebrate. Curr Biol. 16(3):296-300.

73. Vashisht A. A., Zumbrennen K. B., Huang X., Powers D. N., Durazo A., Sun D., Bhaskaran N., Persson A., Uhlen M., Sangfelt O., Spruck C., Leibold E. A., Wohlschlegel J. A. (2009) Control of iron homeostasis by an iron-regulated ubiquitin ligase. Science 326, 718-721.

74. Wang WX, Huang Q, Hu Y, Stromberg AJ, Nelson PT. (2011) Patterns of microRNA expression in normal and early Alzheimer’s disease human temporal cortex: white matter versus gray matter. Acta Neuropathol. 121(2):193–205.

75. Wendler S, Hartmann N, Hoppe B, Englert C. Age-dependent decline in fin regenerative capacity in the short-lived fish Nothobranchius furzeri. (2015) Aging Cell. 14(5):5-857.

76. Wong BX, Duce JA. (2014) The iron regulatory capability of the major protein participants in prevalent neurodegenerative disorders. Front Pharmacol. 5:81.

77. Xu Q. (1999). Microinjection into zebrafish embryos. Methods in Molecular Biology (Clifton, N.J.), 127, 125–132.

78. Yamamoto A., Shin R.W., Hasegawa K., Naiki H., Sato H., Yoshimasu F., and Kitamoto, T. (2002) Iron (III) induces aggregation of hyperphosphorylated tau and its reduction to iron (II) reverses the aggregation: implications in the formation of neurofibrillary tangles of Alzheimer’s disease. J. Neurochem. 82,1137-1147.

79. Zhang DL, Ghosh MC, Rouault TA. (2014) The physiological functions of iron regulatory proteins in iron homeostasis - an update. Front Pharmacol. 13;5:124.

80. Zhang B., Kirov S., & Snoddy J. (2005). WebGestalt: An integrated system for exploring gene sets in various biological contexts. Nucleic Acids Research, 33(SUPPL. 2).

